# Conformational Ensembles of the Disordered 4E-BP2:eIF4E Complex Restrained by smFRET Experiments

**DOI:** 10.64898/2026.04.21.719986

**Authors:** Spencer Smyth, Zi Hao Liu, Thomas E. Tsangaris, Teresa Head-Gordon, Julie D. Forman-Kay, Claudiu C. Gradinaru

## Abstract

Eukaryotic cap-dependent translation initiation is regulated by binding of the predominantly folded eukaryotic initiation factor 4E (eIF4E) to the intrinsically disordered eIF4E binding proteins (4E-BPs). Here, we report full-length atomistic conformational ensembles generated by IDPConformerGenerator and optimized by X-EISDv2 workflow for both apo 4E-BP2, the neuronal 4E-BP, and 4E-BP2 in complex with eIF4E, using data from single-molecule fluorescence and nuclear magnetic resonance (NMR), together with select coordinates from a 4E-BP1:eIF4E crystal structure. Structural sampling within dynamic complexes is often under-appreciated, with NMR and crystal structure data for 4E-BP:eIF4E suggesting different degrees of structural heterogeneity. Our ensemble models validated by solution spectroscopy data enable comparison of free 4E-BP2 and its complex with eIF4E. This shows a delocalization of contacts around canonical regions, which supports previous findings of unidirectional conditional occupancy of the binding sites. Two new contact regions emerged: one between the disordered *N*-termini of eIF4E and 4E-BP2, which may play an allosteric role in tuning the binding affinity, and the other between the C-terminus of 4E-BP2 and an extended region of eIF4E, which is consistent with the extended, dynamic binding interface that we reported previously. These results support a model of translation regulation in which the dynamic 4E-BP2:eIF4E complex facilitates accessibility of regulatory sites of 4E-BP2 when bound.

## INTRODUCTION

Atomic resolution 3D structural information is considered key to delineating the biological function of proteins. Defying this structure-function paradigm, intrinsically disordered proteins and protein regions (IDPs and IDRs hereafter) fail to adopt a stable structure but fulfil many, often regulatory, biological roles, often due to their extreme conformational plasticity (Holehouse and Kragelund 2024). Bioinformatics analysis of the human proteome predicts that about one third of amino acids are disordered, with ∼60% of proteins containing stretches of greater than 30 residues of intrinsic disorder and 5% being completely disordered (Tsang, Pritišanac et al. 2020). IDPs and IDRs are significantly involved in neurological diseases, signalling disorders, diabetes and cancers (Uversky, Oldfield et al. 2008), and developing pharmaceutical drugs that target their pathologies remain a major challenge (Uversky 2025).

The mechanisms by which IDPs/IDRs bind to their molecular partners have generated considerable interest, particularly as potential therapeutic targets (Dogan, Gianni et al. 2014, Csizmok, Follis et al. 2016, Mollica, Bessa et al. 2016, Chen and Kriwacki 2018, Fuxreiter 2019). Local ordering of disordered segments in IDPs/IDRs can be observed upon binding to their biological targets, and in some cases significant folding occurs (Wright and Dyson 2009). In contrast, highly dynamic complexes arise when only a limited number of residues become ordered upon binding and the rest of the protein chain remains flexible, leading to dynamic or “fuzzy” complexes (Sharma, Raduly et al. 2015). Analysis of the correspondence between predicted disordered regions and high confidence structure prediction in AlphaFold2 (Alderson, Pritišanac et al. 2023) leads to an estimate that 15 to 25% of human IDRs conditionally fold upon binding, implying that the majority of the disordered proteome functions in the absence of stable structure, with interactions in the context of dynamic complexes. While AlphaFold-Multimer can reasonably predict structures of stable complexes involving conditionally folding IDRs, predictions fail for highly heterogeneous, fuzzy interactions (Omidi, Møller et al. 2024).

Constructing IDP/IDR conformational ensembles using ensemble-averaged experimental data as restraints is a heavily underdetermined inverse problem (Ravera, Sgheri et al. 2016). Even more challenging is the use of experimental data to refine conformational ensembles of fuzzy complexes of IDRs; for atomistic ensembles of dynamic complexes, the literature largely reports using molecular dynamics (MD) simulations. Although MD can provide insightful information on inter-/intra-molecular interactions, it is not restrained by experimental data and may not achieve sufficient sampling diversity. Here we utilize IDPConformerGenerator (Teixeira, Liu et al. 2022), with its local disorder region sampling (LDRS) approach (Liu, Teixeira et al. 2023), to rapidly model diverse conformations, incorporating NMR chemical shifts to bias the torsion angle distribution of the disordered 4E-BP2 in complex with eIF4E for a high quality data-driven initial pool. Conformers within this pool are then selected based on experimental data using the new X-EISDv2 workflow in the Bayesian X-EISD method (Lincoff, Haghighatlari et al. 2020). Although other reweighting and subsampling methods exist and can be used for dynamic complexes (Bottaro, Bengtsen et al. 2020), X-EISDv2 was chosen for its statistical scoring, speed and highly parameterizable capabilities.

Different experimental observables are sensitive to different properties and length scales of the protein and, when used as restraints, may result in improvement or deterioration of other restraint types, while the impact of different types of restraints may vary for different systems. smFRET has been used to study the binding of IDPs and to compare their conformational and dynamic properties in the free and bound states (LeBlanc, Kulkarni et al. 2018, Metskas and Rhoades 2020, Chowdhury, Nettels et al. 2023). Typically, several labelling positions with different residue separations are selected throughout the sequence to provide sufficient sampling(Borgia, Borgia et al. 2018, Wiggers, Wohl et al. 2021). In particular, intermolecular smFRET is key in defining the overall 3D architecture and topology of the complex. When combined with MD simulations or static conformational ensembles, smFRET data is very efficient to restrain or validate molecular models (Borgia, Borgia et al. 2018, Naudi-Fabra, Tengo et al. 2021, Wiggers, Wohl et al. 2021, Tsangaris, Smyth et al. 2023). A study by Naudi-Fabra et al.(Naudi-Fabra, Tengo et al. 2021) found that six FRET restraints, probing a variety of distances and regions in a 110-residue IDP, were sufficient to restrain the conformational ensemble such that is predictive of FRET data not used as restraints, supporting our use of a significant number of smFRET restraints in this study.

Translation of mRNA to protein is regulated by the tight interaction of eukaryotic initiation factor 4E (eIF4E) with disordered eIF4E binding proteins (4E-BPs) in a phosphorylation-dependent manner (Sonenberg and Hinnebusch 2009). The 120-residue disordered protein 4E-BP2 is found in the brain and regulates synaptic plasticity, which is essential for learning and memory (Banko, Merhav et al. 2007). The 4E-BP2:eIF4E interaction has a 3 nM binding affinity (Lukhele, Bah et al. 2013), yet NMR data revealed that 4E-BP2 binds to eIF4E via a dynamic bipartite interface. This interface includes a canonical helix as primary site (^54^YDRKFLLDRR^63^) and a secondary site (^78^IPGTV^82^), with evidence for interactions spanning residues 34 to 90 (Lukhele, Bah et al. 2013). These results demonstrating an extensive dynamic complex provide additional insights over the crystal structure of the highly similar 4E-BP1:eIF4E complex, which has density only for residues 50-83 of 4E-BP1(Peter, Igreja et al. 2015), but a full structural characterization of the dynamic complex has not been defined.

The ^15^RAIP^18^ motif and the TOR signaling (TOS) motif ^116^FEMDI^120^ facilitate the hierarchical phosphorylation of 4E-BPs by the mammalian target of rapamycin (mTOR) kinase (Böhm, Imseng et al. 2021). Five-site phosphorylation of 4E-BP2 reduces its binding affinity by nearly four orders of magnitude by inducing a β-sheet folded domain that sequesters the primary binding motif (Bah, Vernon et al. 2015), see **Fig. 1**. Two of the phosphorylation sites are in the folded domain and 3 are in the C-terminal IDR (C-IDR). The C-IDR phospho-sites modulate the folded domain stability, which effectively tunes access to eIF4E-binding sites via changes in conformational equilibria (Dawson, Bah et al. 2020).

**FIGURE 1.**
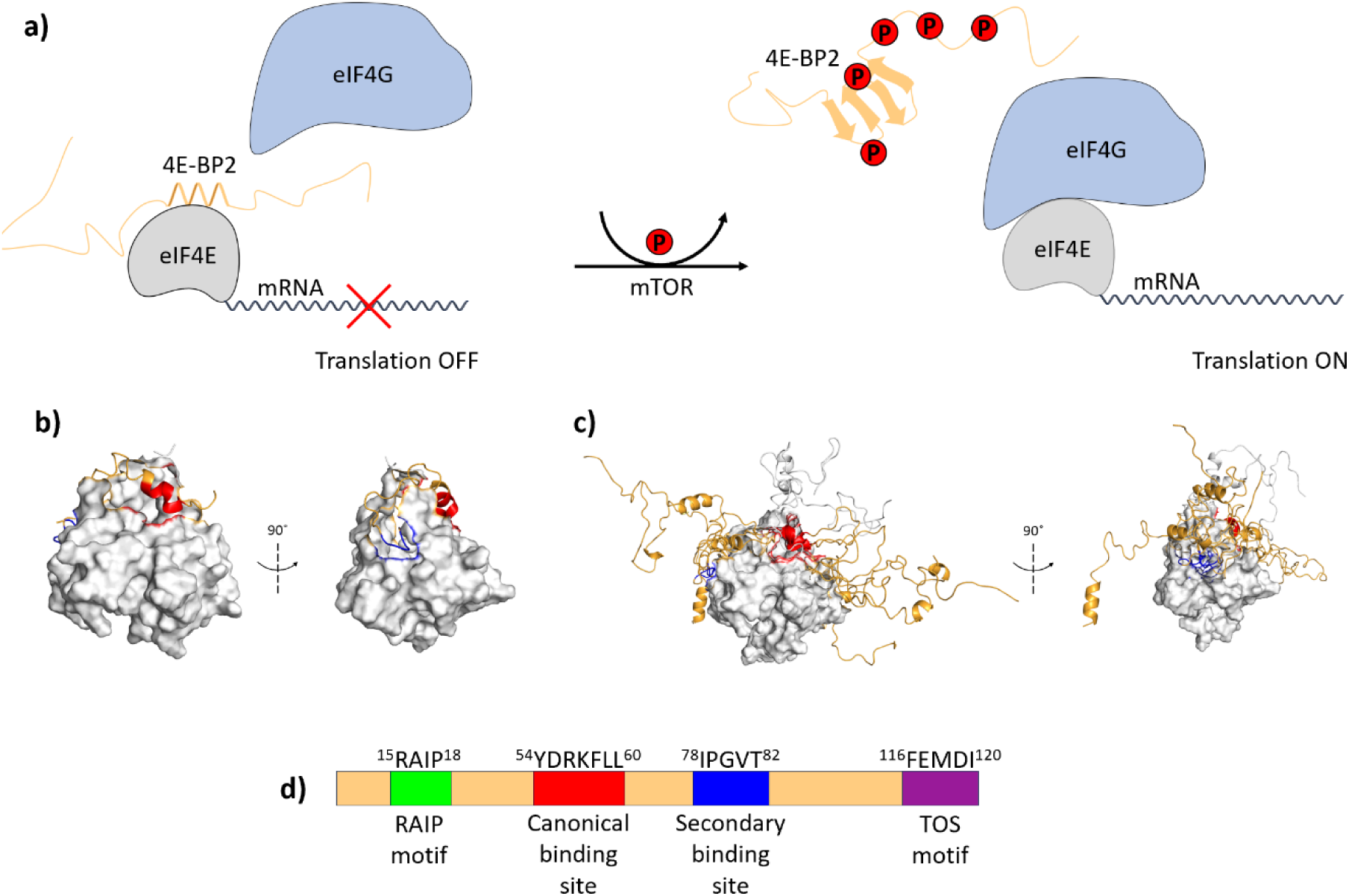
Schematic overview of the biological role and of the 4E-BP2 and eIF4E proteins in cap-dependent translation. Key sequence features and structural motifs of the 4E-BP2:eIF4E protein complex are shown. (**a**) Cap-dependent translation is mediated by eIF4E binding to mRNA at the 5’ -cap, 4E-BP2 inhibits the initiation of translation by competing with eIF4G to bind the shared binding site on eIF4E. Phosphorylation by mTOR reduces the binding affinity of 4E-BP2 allowing eIF4G to outcompete it form a translation competent complex by association with other proteins. (**b**) Structure of the complex of eIF4E complex with a fragment of 4E-BP1 (yielding coordinates for residues 50-83) (PBD ID: 4UED). (**c**) Conformational ensemble derived from the 4UED crystal structure and IDPConformerGenerator. (**d**) Schematic of the 4E-BP2 sequence with the important features colored: RAIP motif, canonical binding motif (CBM), secondary binding motif (SBM), and the mTOR signaling (TOS) motif.

In our previous work, smFRET measurements were used to delineated the flexible and extended nature of eIF4E-bound 4E-BP2 conformations and revealed that the C-terminal region also contributes to binding interactions (Smyth, Zhang et al. 2022). However, a detailed structural picture of the complete 4E-BP2:eIF4E complex, reflecting its heterogeneous and dynamic ensemble, has not been described. Here, we report a conformational ensemble of the 4E-BP2:eIF4E complex, built by IDPConformerGenerator (Teixeira, Liu et al. 2022) starting from the X-ray crystal structure template of the 4E-BP1:eIF4E complex (Peter, Igreja et al. 2015), with extensive sampling of dynamic elements (based on NMR data) restrained by data from eight intramolecular and two intermolecular FRET pairs (**Fig. 1 b-c**) using X-EISDv2.

The resulting ensemble, consistent with experiment, has significant structural heterogeneity not accurately represented in the crystal structure. It demonstrates contacts between the C-terminus of 4E-BP2 and both binding sites on eIF4E from the crystal structure, corroborating previous observations of binding-induced changes of this region of 4E-BP2 (Smyth, Zhang et al. 2022). The ensemble also showed significant intramolecular eIF4E contacts between a region of the N-IDR that was reported to act as a 4E-BP2 binding inhibitor and the canonical binding site (Abiko, Tomoo et al. 2007). The results highlight the importance of carefully considering both the method of prior ensemble generation and the type of experimental data in the modelling of IDP/folded protein dynamic complexes. This realistic and experimentally restrained conformational ensemble of the 4E-BP2:eIF4E complex advances our understanding of translational regulation and provides new insights into how dynamic complexes of disordered proteins facilitate function.

## RESULTS

Generation of realistic conformational ensembles of IDPs in dynamic complexes is challenging due to a massive sampling space, a lack of PDB learning data for efficient generation of physical priors, and a scarcity of biophysical data acting as hard optimization restraints. Comparative analysis of the target-bound *vs.* the apo IDP ensembles is a powerful tool to identify key intra- and inter-molecular contacts that drive the equilibrium and illuminate potential modulatory and pharmacological pathways. Based on this, we first refined the conformational ensemble of 4E-BP2 in the free state before modelling the eIF4E-bound ensemble.

### Multiple smFRET Constructs Probe the Heterogeneity of 4E-BP2

Previous smFRET measurements (Smyth, Zhang et al. 2022) revealed different degrees of expansion and stiffening for two segments of the 4E-BP2 chain (32-91 and 73-121) upon binding to eIF4E. A follow-up computational study (Tsangaris, Smyth et al. 2023) showed that FRET efficiencies were the most powerful restraints for the refinement of non-phospho and 5-phospho 4E-BP2 conformational ensembles. Motivated by these findings and other recent studies (Naudi-Fabra, Tengo et al. 2021, Wiggers, Wohl et al. 2021), six additional FRET constructs were designed to systematically probe the entire 4E-BP2 protein chain. The sequence was split into three labelling segments: 0-32, 32-91 and 91-121, hereafter denoted as segments ***A***, ***B*** and ***C***, respectively. To provide richer complementary topological information, all additional two-residue combinations of residues 0, 32, 91 and 121 were sampled resulting in constructs ***AB*** (0-91), ***BC*** (32-121), and ***ABC*** (0-121). This recapitulates a previous approach (Wiggers, Wohl et al. 2021) to probe the dynamics of the cell adhesion protein E-cadherin as it binds to its folded binding partner β-catenin. An additional construct with labelling positions at residues 14 and 73 (***β-fold***) was used to probe the chain segment that undergoes folding to a four-stranded β-sheet in the five-fold phosphorylated (5P) state; all FRET constructs are summarized in **Fig. S1**.

As shown in **Fig. 2**, diffusion-based smFRET measurements were performed on all six 4E-BP2 double-cysteine mutants; for completeness, smFRET data for the ***B*** (H32C/S91C) and ***C-term*** (C73/C121) constructs published previously (Smyth, Zhang et al. 2022) are also included. FRET efficiency generally decreases with the number of residues in a segment, *N*_*r*_, i.e., the sequence separation between labels, with some notable exceptions. The ***AB*** and ***BC*** segments have similar *N*_*r*_ but their 〈*E*〉 values differ substantially, while segments ***B*** and ***β-fold*** exhibit the highest FRET despite not having the lowest *N*_*r*_. These differences are attributed to 4E-BP2’s unique intramolecular interactions. The protein is overall more compact than a statistical coil, as the measured FRET efficiencies for all segments are either higher or identical, within uncertainty, to those predicted for a statistical coil (**Table S1**).

**FIGURE 2.**
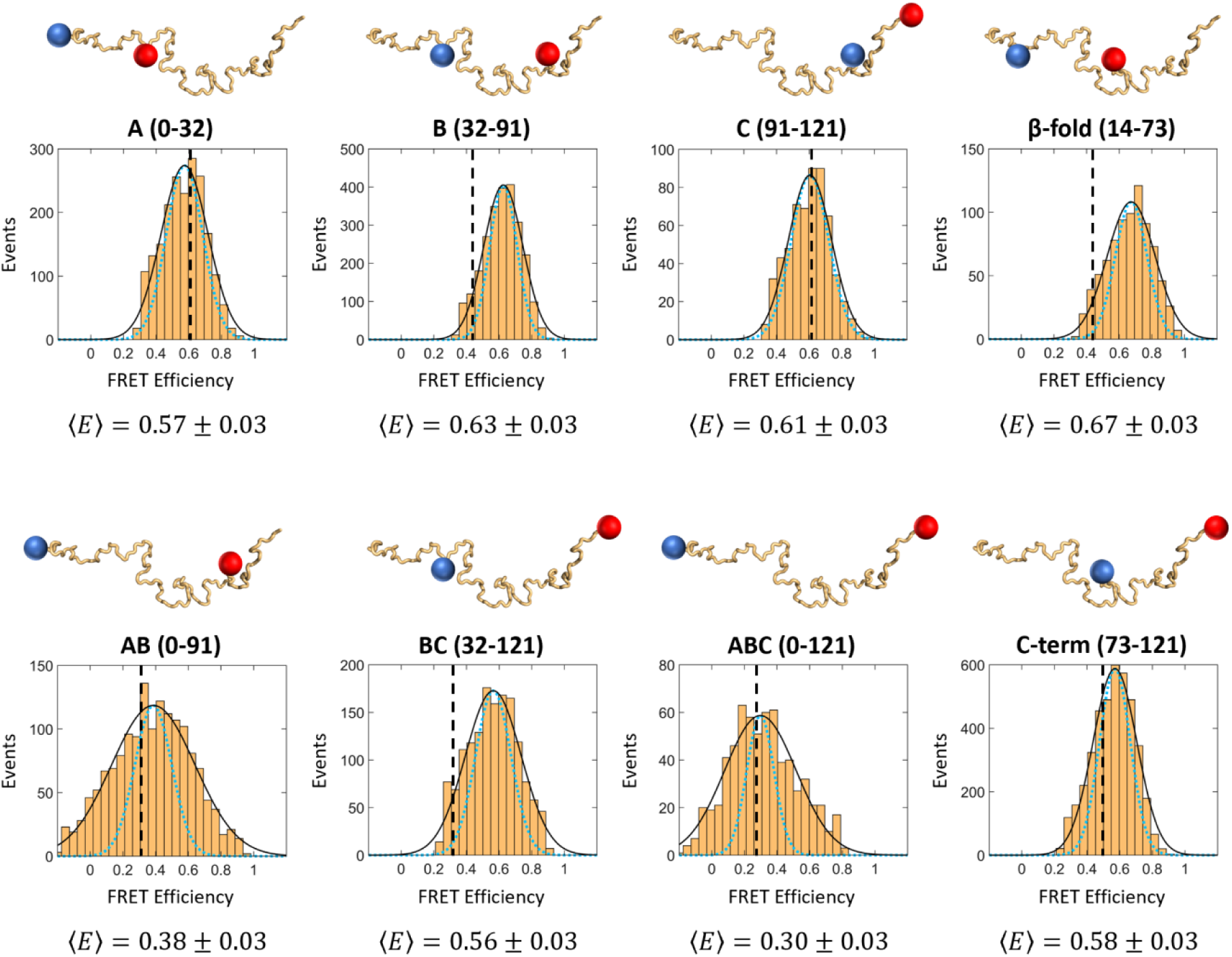
Intramolecular smFRET measurements on eight complementary segments of 4E-BP2: *A, B, C, β-fold, AB, BC, ABC, and C-term*. The structures of 4E-BP2 labelled with red and blue spheres indicate the labelling positions of the donor and acceptor fluorophores, respectively. Each FRET histogram was fitted to a Gaussian distribution (black line), with the mean FRET efficiency 〈*E*〉 ± s.e.m. (standard error of the mean) listed under each panel. Shot-noise limited (SNL) distributions for each dataset (blue dashed line) are normalized Gaussian distributions calculated as described in SI. Vertical dashed black lines indicate the mean FRET efficiency of each segment predicted from the TraDES random coil ensemble (see Methods). Data for B (H32C/S91C) and C-term (C73/C121) were previously published (Smyth, Zhang et al. 2022).

The excess width of the experimental FRET distributions compared to the shot-noise limit (*R*_σ_ = σ_*obs*_⁄σ_*sn*_) shows a positive correlation with the length of the segment, *N*_*r*_(**Fig. S2a**). *R*_σ_ could be caused by internal chain dynamics on time scales longer than the average inter-photon time, 1-10 μs, but shorter than the typical burst duration (∼1 ms) in our experiments (Gopich and Szabo 2012). In the Rouse model (Rouse Jr 1953), the Rouse time (τ_*R*_) is the relaxation time of the lowest mode and τ_*R*_ ∝ *N*_*r*_^2^; it is thus expected that the slower chain reconfiguration time will lead to a broadening of the FRET histogram at larger *N*_*r*_.

### An Improved 4E-BP2 Conformational Ensemble

Considering these new data, we evaluated the predictive ability of our previously published 4E-BP2 ensemble (Tsangaris, Smyth et al. 2023), which was restrained with SAXS, NMR chemical shifts and two FRET efficiencies (***B*** & ***C-term***), see **Table S2**. Four of the six new FRET values are predicted by the previous model within the uncertainty. This is in agreement with the expectation from Naudi-Fabra et al.(Naudi-Fabra, Tengo et al. 2021), where fewer than six FRET restraints was insufficient to be predictive of additional FRET efficiencies for a 110 residue chain. The ***BC*** and ***β-fold*** segments are more compact than predicted from the previous ensemble as indicated by their higher measured FRET efficiencies. In light of these results, the previously identified long-range contacts between residues ∼20-40 and ∼80-100 (Tsangaris, Smyth et al. 2023) likely include the C-terminal end residues (100-120), as well.

Here, a much larger initial pool of conformers (10^5^ *vs* 2×10^4^) was generated by IDPConformerGenerator (Teixeira, Liu et al. 2022) compared to FastFloppyTail (FFT) (Ferrie and Petersson 2020) used previously (Tsangaris, Smyth et al. 2023). In addition, experimentally derived NMR *C_α_* and *C_β_* chemical shifts (Lukhele, Bah et al. 2013, Dawson, Bah et al. 2020) were used to bias the secondary structural content in IDPConformerGenerator. From this initial pool, the Bayesian X-EISD method (Lincoff, Haghighatlari et al. 2020), using the newly developed X-EISDv2 (see Methods), was used to select a subset of 100 conformers that are simultaneously in agreement with paramagnetic relaxation enhancement (PRE) data (Tsangaris, Smyth et al. 2023), the hydrodynamic radius (*R*_ℎ_) measured by FCS (Smyth, Zhang et al. 2022), and all eight smFRET 〈*E*〉 values reported here. During optimization, the FRET data were given more weight to achieve reasonable agreement with all eight restraints; a weight factor of Ω = 50 was used which is similar to Ω = 40 used previously (Tsangaris, Smyth et al. 2023). The normalized chi-squared 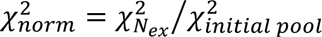 for each restraint was used to determine the optimum number of conformer exchanges (*N*_*ex*_), guided by the knee-point of the optimization curve (**Fig. S3**). At *N*_*ex*_ = 10^5^, good agreement was achieved with PRE (**Fig. S4**), *R*_ℎ_ (**Table S3**), and FRET (**Fig. S5**), and this optimized ensemble was used for all subsequent analysis.

The internal scaling profile (ISP) of the optimized ensemble (**Fig. 3a**) represents the mean inter-residue distances (*R*_|*i*−*j*|_) as a function of residue separation (|*i* − *j*|). In the “*short*” range, 1 ≤ |*i* − *j*| ≤ 30, the 4E-BP2 scaling resembles that of the TraDES random coil (RC); however, in the “*intermediate*” range, 30 ≤ |*i* − *j*| ≤ 100, the scaling exponent (*v*) decreases steeply. The minimum value of *vv* is at |*i* − *j*|∼70, corresponding to a change in concavity of the ISP curve (**Fig. S6**), whereas in the “*long*” range, 100 ≤ |*i* − *j*| ≤ 120, *v* > 0.5 and it monotonically increases. This stark deviation from uniform scaling behavior is typically attributed to different types of inter-residue interactions; in the case of 4E-B2, attractive in the intermediate range and repulsive in the long range. As previously reported (Tsangaris, Smyth et al. 2023), the significant 4E-BP2 chain compaction at the intermediate inter-residue range can be explained by invoking net-charge-per-residue (NCPR) arguments. Three oppositely charged regions separated by ∼70 residues in the 4E-BP2 sequence likely drive this compaction: 11-24 (+ NCPR) with 85-98 (- NCPR), 22-37 (- NCPR) with 103-111 (+ NCPR), and 47-63 (+ NCPR) with 108-121 (- NCPR).

**FIGURE 3.**
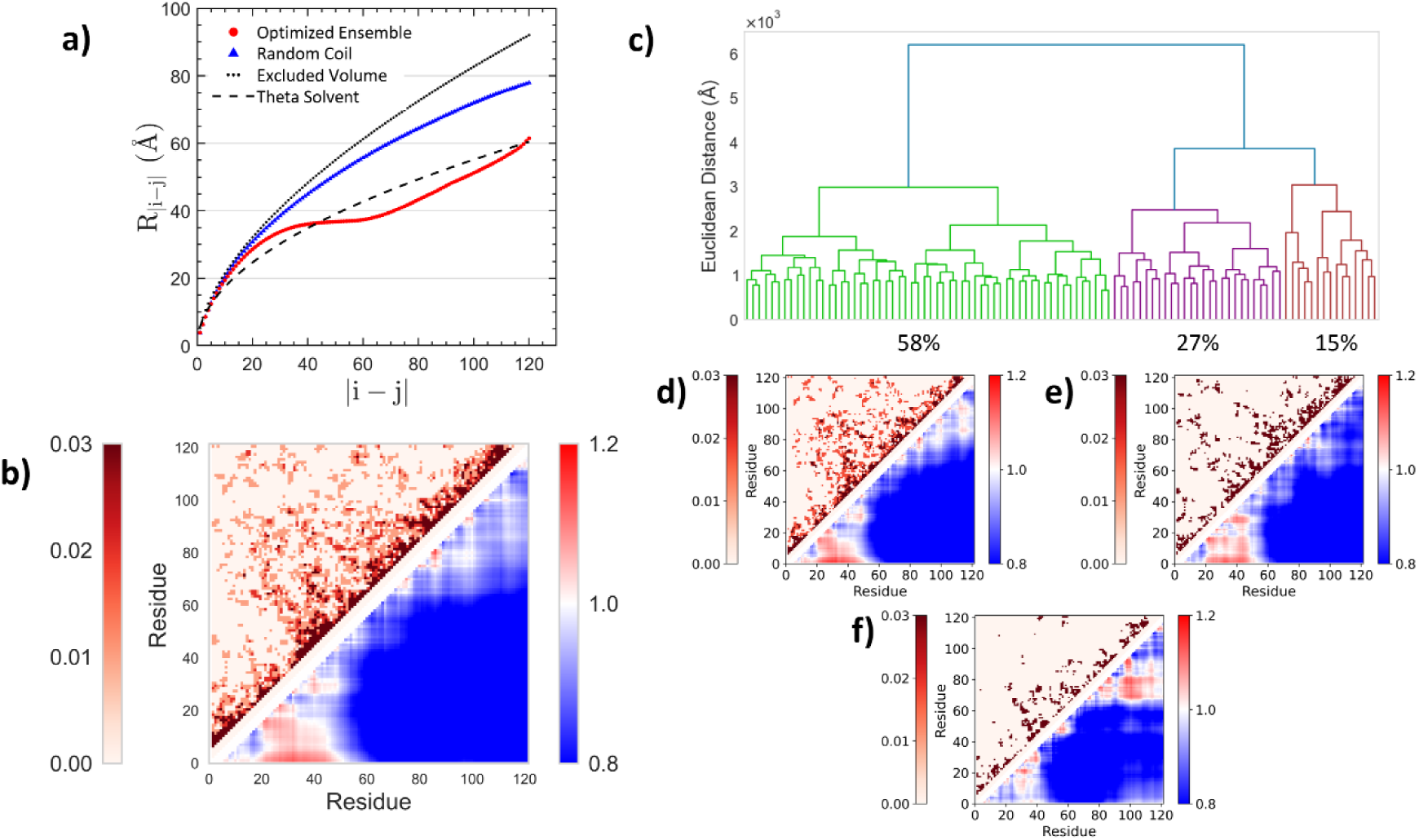
Analysis of the optimized conformational ensemble of 4E-BP2. (**a**) Internal scaling profiles of the optimized ensemble (red), TraDES random coil (RC) ensemble (blue), excluded-volume, EV (black, dotted), and *θ*-solvent (black, dashed) homopolymer models. (**b**) *Upper*: contact map showing the fraction of inter-residue contacts in the optimized ensemble; two residues are in contact if their *C_α_* atoms are less than 8 Å apart. *Lower*: distance map of the optimized ensemble normalized by the TraDES random coil ensemble. (**c**) Dendrogram showing the agglomerative hierarchical clustering of the optimized 4E-BP2 ensemble: *Cluster 1* (green), *Cluster 2* (purple), and *Cluster 3* (brown). Contact maps and inter-residue distance maps normalized by the TraDES RC ensemble for clusters 1-3 (**d**-**f**).

Two-dimensional maps of contact frequency and inter-residue distance are typically used to delineate interaction patterns in a conformational ensemble (**Fig. 3b**). These maps are constructed based on the statistics of inter-residue *C_α_ – C_α_* distances: the distance map was normalized by corresponding values in the random-coil ensemble, and two residues are defined as being in contact if the *C_α_ – C_α_* distances < 8 Å. The normalized 2D inter-residue distance map shows significant compaction compared between residues ∼1-60 and ∼60-121, this extends the region of residues ∼20-40 and ∼80-100 identified in the previous optimized 4E-BP2 ensemble (Tsangaris, Smyth et al. 2023). All three of the opposite NCPR regions now fall within the observed range of compaction, highlighting the importance of the additional FRET restraints which are likely to drive this. Contacts are well distributed throughout the sequence with the highest density occurring between residues ∼20-40 and ∼60-80.

Agglomerative hierarchical clustering was applied to the optimized ensemble using the Euclidean distance between inter-residue distances in different conformers as a dissimilarity metric (Baul, Chakraborty et al. 2019, Tsangaris, Smyth et al. 2023). The dendrogram split into a total of four clusters (**Fig. S7**) before the cutoff criterion was satisfied (**Fig. S8a**); due to the low population (∼5%) *Cluster 3* was combined with *Cluster 4* resulting in three final clusters (**Fig. 3c**). As shown in **Fig. 3 d-f**, *Clusters 1* and *2* show similar patterns of inter-residue distances as the complete optimized ensemble with the most dissimilar being *Cluster 3*, comprising 15% of the optimized ensemble. *Cluster 3* has significant expansion between residues (65-85) and (85-115) and a reduced region of expansion in the N-terminal region, while also having the lowest number of intramolecular contacts. *Clusters 2* exhibits the largest number of contacts, while in *Cluster 1* they are largely dispersed throughout the sequence. Despite being most prominent in *Clusters 1* and *2*, all clusters show an expansion around residue T37, reflecting accessibility and consistent with T37 being the first site phosphorylated in the hierarchical order (Gingras, Gygi et al. 1999).

### smFRET on the 4E-BP2:eIF4E complex

To generate solution-based restraints for the 4E-BP2:eIF4E dynamic complex, smFRET histograms of all eight 4E-BP2 segments were also measured in the presence of ∼10^4^ excess (2 µM) eIF4E (**Fig. 4a**). Based on a dissociation constant *K*_*d*_ = 3.2 *nM* (Bah, Vernon et al. 2015), more than 99% of 4E-BP2 molecules are bound to eIF4E. Changes in FRET efficiency upon binding to eIF4E are heterogeneous across the eight intramolecular segments measured. For five segments (***B***, ***AB***, ***BC***, ***β-fold***, and ***C-term***) the FRET decreases indicating an upshift of their distance distributions, two segments (***A*** and ***C***) show an increase in FRET efficiency indicating a downshift, while for one segment (***ABC***) the FRET histogram remains unchanged. A similar combination of expansion and compaction of different IDP segments upon interaction with a folded target was observed in the dynamic complex between the disordered E-cadherin and the folded β-catenin (Wiggers, Wohl et al. 2021). There, five segments of E-cadherin compacted to varying degrees in the bound state, while part of the N-terminal region of E-cadherin expanded; this was attributed to the disruption of the electrostatic interactions of the opposite charges of this segment in the free state.

**FIGURE 4.**
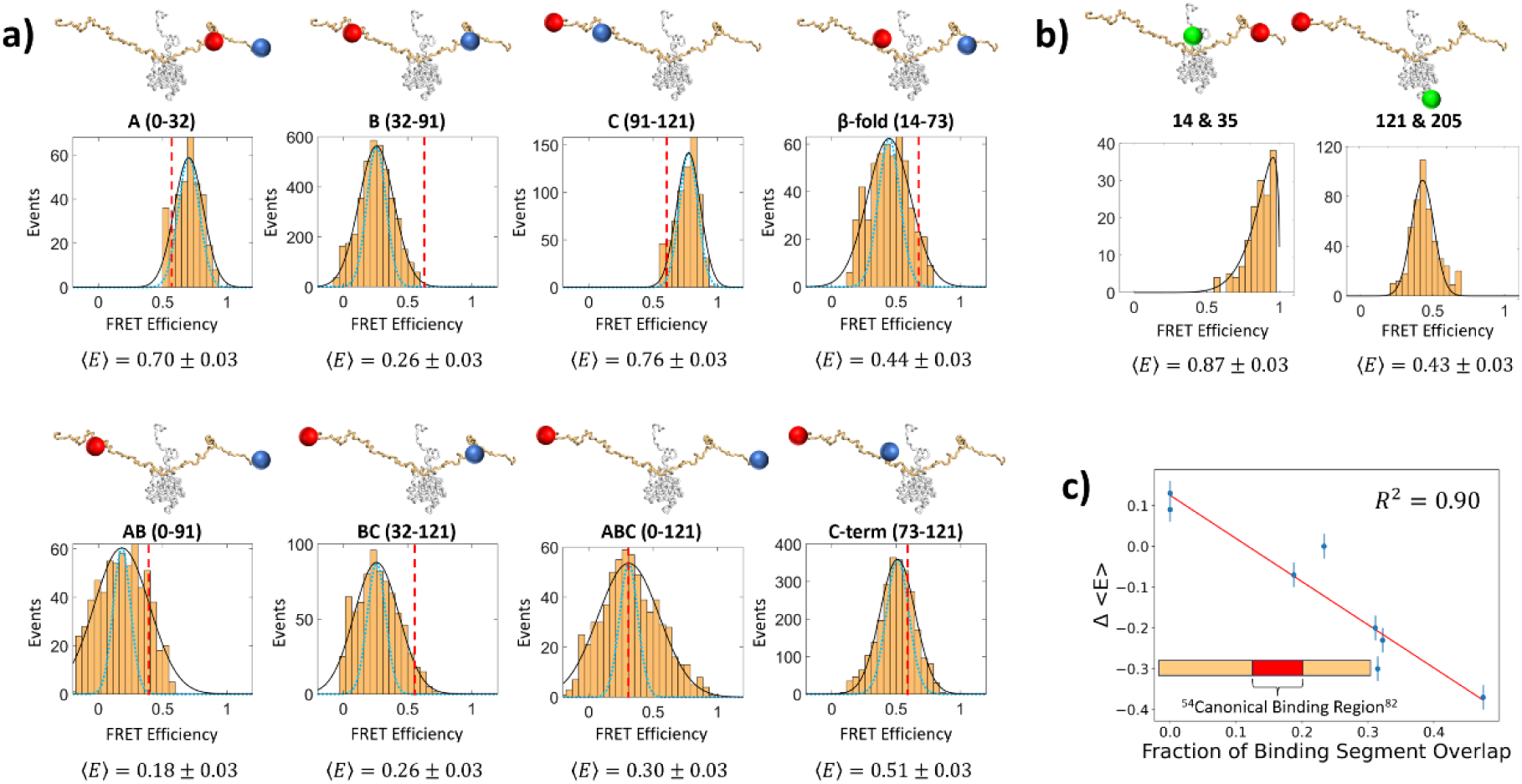
(**a**) smFRET histograms of eight segments of 4E-BP2 bound to eIF4E: *A, B, C, β-fold, AB, BC, ABC,* and *C-term*. The structures of 4E-BP2 labelled with red and blue sphere indicate the labelling positions of the donor and acceptor fluorophores, respectively. FRET histograms were each fitted to a Gaussian distribution (black line), with 〈*E*〉 ± s.e.m. listed under each panel. SNL distributions for each dataset (blue dashed line) are normalized Gaussian distributions calculated as described in SI. Vertical dashed red lines indicate 〈*E*〉 of each segment measured in the apo state. (**b**) Intermolecular smFRET histograms with 4E-BP2 labelled with Cy5 at residue 14 or 121 and eIF4E labelled with Cy3 at residue 35 or 205, respectively. Histograms were fitted to a non-normalized Beta distribution and a single Gaussian (solid black lines), respectively, with 〈*E*〉 ± s.e.m. listed under each panel. (**c**) The change in 〈*E*〉 of 4E-BP2 upon binding to eIF4E is plotted against the fraction of each segment that overlaps with the canonical binding region (residues 54-82).

Previous studies have highlighted the utility of intermolecular smFRET in the characterization of disordered proteins in protein-protein complexes and biomolecular condensates (Iljina, Garcia et al. 2016, Borgia, Borgia et al. 2018). Intermolecular FRET provides critical distances between interacting partners that constrain their relative orientations in the complex and complement the intramolecular FRET restraints. To address this need, TIRF-measured smFRET data were obtained on surface-immobilized 4E-BP2:eIF4E particles having a different fluorophore attached to each protein (**Fig. 4b**).

The changes in FRET efficiencies of different 4E-BP2 segments upon binding to eIF4E reflect the degree to which each segment interacts with known binding elements of eIF4E. Changes in FRET efficiency are negatively correlated with the degree to which the segment probed overlaps with the ‘binding region’ of 4E-BP2, which is defined by residues 54-82. This region includes both the canonical and secondary eIF4E-binding sites (Lukhele, Bah et al. 2013), and the residues within this region are expected to experience the largest binding-induced changes. For instance, segment ***B*** has the largest fraction of residues that overlap with the binding region and shows the largest decrease in FRET efficiency, suggesting that this segment forms extended conformers that wrap around the eIF4E scaffold to facilitate interactions of both binding sites. In contrast, segments ***A*** and ***C*** do not overlap at all with the binding region and are the only 4E-BP2 constructs that show an increase in FRET efficiency upon binding. When the change in FRET efficiencies between free and eIF4E-bound states of 4E-BP2 are plotted against the fraction of the segment’s residues that overlap with the binding region (**Fig. 4c**), a strong negative correlation (*R*^2^ = 0.90) is observed. This suggests that the FRET changes can be explained in large part by the extended topology of a multisite interaction model for the 4E-BP2: eIF4E complex. As for the apo state (**Fig. 2**), the strong correlation between the excess width of the FRET distributions *vs.* the shot-noise limit (*R*_σ_ = σ_*obs*_⁄σ_*sn*_) and the segment length, *N*_*r*_(**Fig. S2b**), is a signature of slower intrachain dynamics.

It is also interesting to note that segments ***A*** and ***C***, which do not overlap at all with known binding site residues, compact rather than expand in the complex with eIF4E, suggesting that binding favors a larger number of intramolecular 4E-BP2 contacts compared to the free state. This compaction may have functional implications, as the ***A*** segment contains the ^15^RAIP^18^ motif and the ***C*** segment contains the TOS) motif (^116^FEMDI^120^) (Tee and Proud 2002), both of which are important for the phosphorylation of 4E-BP2 by the mTOR kinase (Nojima, Tokunaga et al. 2003, Schalm, Fingar et al. 2003). mTOR associates with 4E-BP2 through a scaffolding protein, the 150 kDa regulatory-associated protein of mTOR (Raptor), which is known to bind to both the TOS and RAIP motifs of 4E-BP2 (Böhm, Imseng et al. 2021). One might expect that compaction upon binding would decrease the accessibility of these sites to the large Raptor protein. However, another remember of the 4E-BP family, 4E-BP1, binds to Raptor at both sites independently of its association with eIF4E (Böhm, Imseng et al. 2021).

### Generation and Optimization of the eIF4E:4E-BP2 Ensemble

In recent years, an increasing number of IDP structural ensembles have been deposited in the Protein Ensemble Database(Ghafouri, Lazar et al. 2023), although most of these entries are for proteins in their free, unbound states. Calculating ensembles of complexes of disordered proteins bound to their (folded) molecular targets remains a challenging prospect, inhibiting understanding of sequence-ensemble-function relation for IDPs (Hadži, Loris et al. 2021). To model the eIF4E-bound state of 4E-BP2, we made use of existing structural information from a high resolution crystal structure of the 4E-BP1:eIF4E complex (PDB ID: 4UED) (Peter, Igreja et al. 2015). This structure was used because it resolves both canonical and secondary binding sites and because 4E-BP1 and 4E-BP2 have the same eIF4E binding mechanism with high sequence identity for binding residues (Fletcher, McGuire et al. 1998). The overall sequence identity of 4E-BP1 and 4E-BP2 is 57%; for the fragment of 4E-BP1 observed in the crystal structure, the sequence identity is 85%. The crystal structure has coordinates for 4E-BP1 residues 50-83 and most of the eIF4E residues, only lacking the N-terminal disordered tail (residues 1-32).

The initial conformation pool of the 4E-BP2:eIF4E complex was built starting from the X-ray crystal structure (PDB ID: 4UED) (Peter, Igreja et al. 2015), see **Fig. 1b-c**. Coordinates for residues of 4E-BP2 and eIF4E that were not resolved in the crystal structure were generated with IDPConformerGenerator (Teixeira, Liu et al. 2022), using the local disordered region sampling (LDRS) tool (Liu, Teixeira et al. 2023). IDPConformerGenerator samples φ, ψ, and ω torsion angles from the PDB. Given the dynamic nature of 4E-BP2 in the complex (Lukhele, Bah et al. 2013, Smyth, Zhang et al. 2022), it is unlikely that the conformation of residues 50-83 in the crystal structure of the complex is the only one present in solution. As such, three cases were modelled by: **1**) fixing only the canonical binding site (*Canonical*, residues 54-60), **2**) fixing only the secondary binding site (*Secondary*, 78-82), and **3**) fixing both binding sites (*Both*, residues 54-60 and 78-82). **Fig. 5a** shows the assembly process of the 4E-BP2:eIF4E conformational ensemble; starting from the crystal structure all residues that are not fixed in each specific instance are removed, then ensembles of the disordered loops or tails are screened with LDRS to select those consistent with the required geometry. A subset of structures for each of the *Canonical*, *Secondary*, and *Both* ensembles is shown in **Fig. 5b**.

**FIGURE 5.**
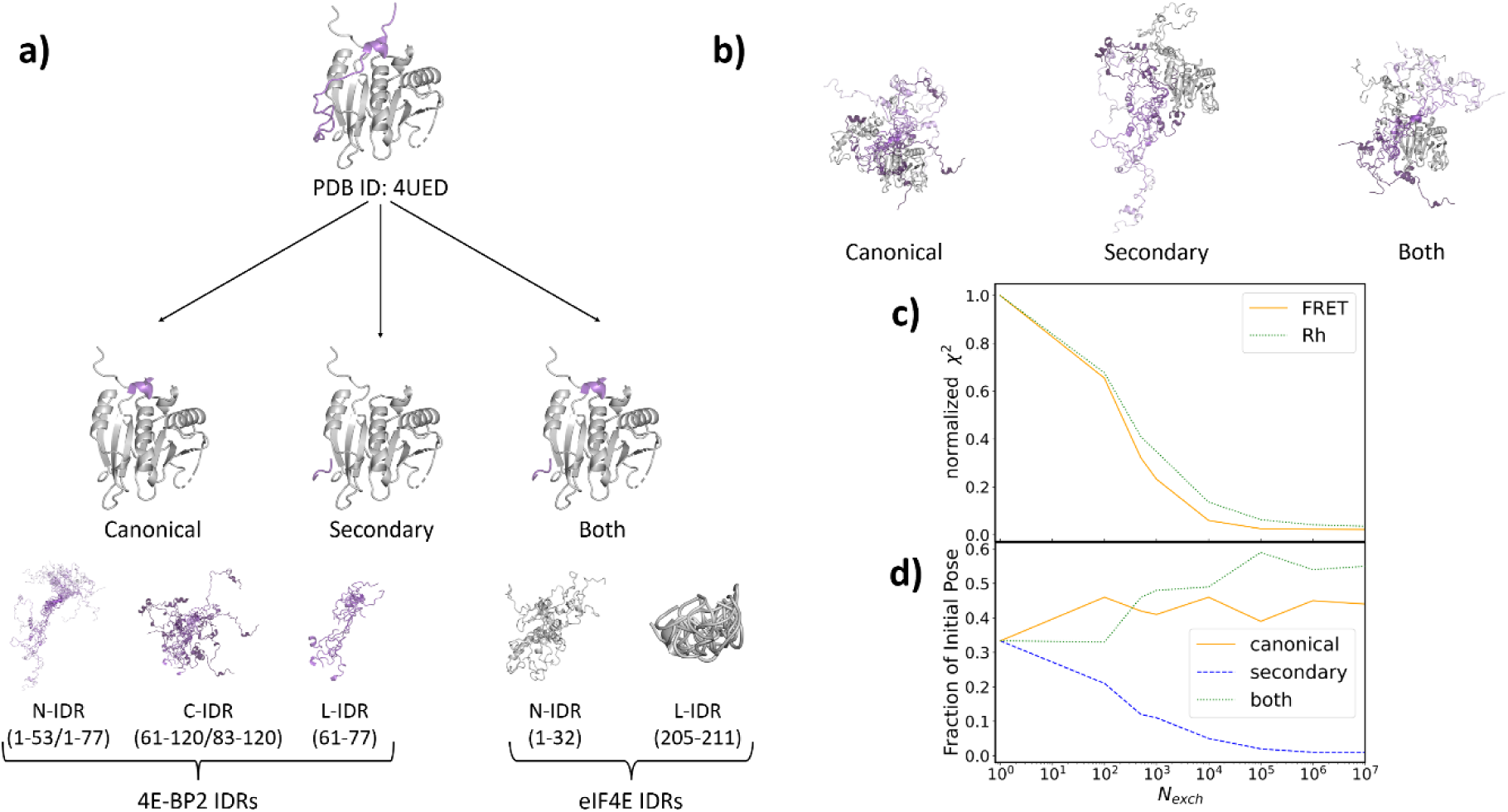
(**a**) Visualization of the construction of the 4E-BP2:eIF4E complex conformational ensemble starting with the PDB ID: 4UED structure. From the initial crystal structure, a subset of residues is selected: for eIF4E the N-IDR (residues 1-32) and L-IDR (residues 205-211) are generated using IDPCG for all structures. For 4E-BP2 three distinct starting pools were used: the *Canonical*, where residues 54-60 were fixed, *Secondary*, where residues 78-82 were fixed, and *Both*, where residues 54-60 and 78-82 were fixed. IDPCG was used to sample all remaining residues. (**b**) Ensembles of five structures are shown for each of the *Canonical, Secondary,* and *Both* cases. (**c**) X-EISD optimization: as the number of conformer exchanges (*N_exch_*) increases, the normalized *χ^2^*between back-calculated and experimental data decreases. (**d**) Contribution of conformers from the three starting pools in the optimized 4E-BP2:eIF4E ensemble as a function of *N_exch_*.

The new X-EISDv2 version of the Bayesian-based method, X-EISD (Lincoff, Haghighatlari et al. 2020), was used to select a 100-conformer subset of the initial pool of structures that are in agreement with all 8 intramolecular FRET efficiencies of 4E-BP2 when bound to eIF4E and the two intermolecular FRET efficiencies (**Fig. S9**), as well as the hydrodynamic radius *R_h_* of the complex (**Table S3**). **Fig. 5c** shows the convergence of the normalized error, 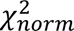, as the number of exchanges *N* increases; as both FRET and *R_h_* share a common knee-point at around *N*_*exch*_ = 10^5^, this ensemble was used for subsequent analysis. **Fig. 5d** shows the fractions of conformers in the restrained ensembles that originate from the *Canonical*, *Secondary*, and *Both* cases as a function of *N*_*exch*_. By design, the initial pool contains equal numbers for each type of conformer, but during refinement the ensemble becomes depleted in *Secondary* conformers, while being enriched in nearly equal amounts of *Canonical* and *Both* conformers. At *N*_*ex*_ = 10^5^ the *Canonical* fraction is 0.39, *Secondary* is 0.02, and *Both* is 0.59. Interestingly, when all 4E-BP2 residues in the crystal structure were fixed in the initial pool (*Both*), no subset of conformations that agreed with all restraints could be found (Smyth 2024). In contrast, starting from the mixed pool containing *Canonical*, *Secondary*, and *Both* ensembles, subsets of conformations that satisfied all experimental restraints were found. This points to additional structural heterogeneity of the 4E-BP2:eIF4E complex which is present in solution (Lukhele, Bah et al. 2013), but is not accurately represented in the crystal structure.

Previous binding studies show that peptide fragments containing only the canonical binding site bind with micromolar affinity, while no binding was detected for the H74-E89 fragment containing only the secondary binding site (Paku, Umenaga et al. 2012). Furthermore, the interaction of the canonical site appears to be enthalpically driven, with the secondary site entropically driven (Lukhele, Bah et al. 2013). As such, given the dynamic nature of the secondary binding site, the fraction of conformers where the secondary binding site is fixed to the crystal structure geometry is expected to be low, consistent with our optimized ensemble.

### Binding-induced Changes in the 4E-BP2 Ensemble

2D inter-residue contact difference and normalized distance maps were constructed to compare the optimized ensembles of 4E-BP2 in the bound and free states (**Fig. 6a**). 4E-BP2 when bound to eIF4E is expanded overall compared to the apo state, especially between residues 1-60 with 60-120 which reverses the contraction of these separations seen in **Fig. 3a**. Topologically, more extended conformations of 4E-BP2 that wrap around eIF4E enable it to interact with the extended binding interface on the surface of eIF4E(Lukhele, Bah et al. 2013). At the same time, there is compaction compared to the apo state observed for *N*- and *C*-terminal stretches of ∼20-30 residues. Thus, the phospo-regulatory ^15^RAIP^18^ motif is brought closer to the first two phosphorylation sites T37 and T46, potentially pre-forming conformations that facilitate the initial phosphorylation steps.

**FIGURE 6.**
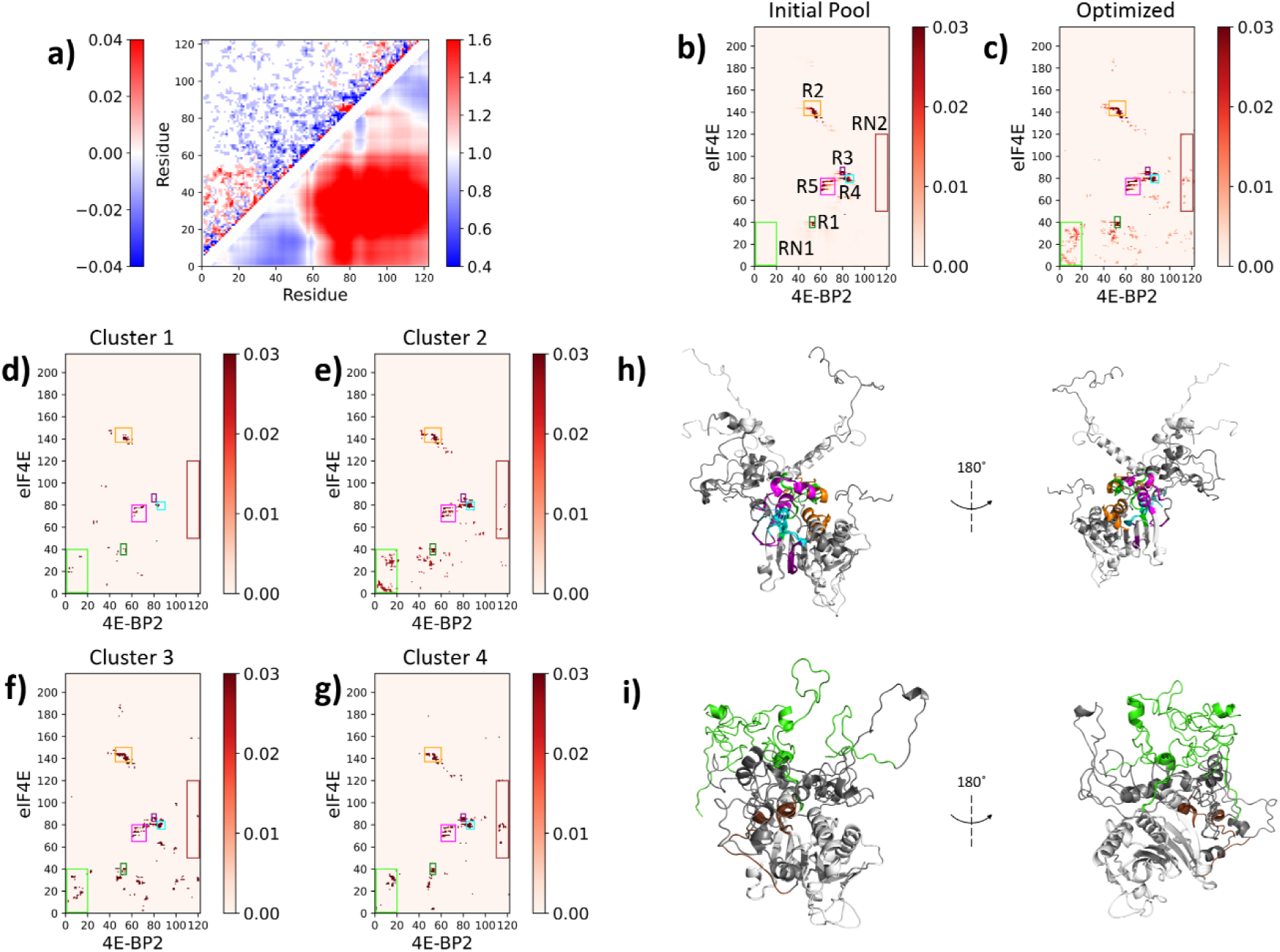
Analysis of the optimized conformational ensemble of the 4E-BP2:eIF4E complex. (**a**) Intramolecular difference contact map (upper) and normalized distance map (lower) of the 4E-BP2 protein in the bound vs the apo state; residues are in contact if their *C_α_* atoms less than 8 Å apart. Intermolecular contacts map between 4E-BP2 (*x-axis*) and eIF4E (*y-axis*) residues in the initial pool ensemble (**b**) and the optimized ensemble (**c**); colored boxes outline regions of significant intermolecular contacts in the initial pool (R1-R5) and emergent contacts in the optimized ensemble (RN1, RN2). (**d-g**) The 4E-BP2:eIF4E contact maps of clusters 1-4 resulted from hierarchical clustering of the optimized ensemble (**Fig. S10**) (**h**) Conformer(s) from the 4E-BP2:eIF4E initial pool selected to illustrate R1-R5 contacts using the same color scheme as in (**b-g**). (**i**) Conformer(s) from the 4E-BP2:eIF4E optimized ensemble selected to illustrate the emergent RN1 & RN2 contacts identified in (**c**).

### Intermolecular Contacts in the 4E-BP2:eIF4E Ensemble

2D inter-residue contact maps for the initial pool (**Fig. 6b**) and the optimized (**Fig. 6c**) 4E-BP2:eIF4E ensembles were constructed for all intermolecular residue combinations. Five regions (*R1-R5*) of prominent intermolecular contacts were identified in the initial pool: *(R1)* residues ∼50-55 of 4E-BP2 with residues ∼35-45 of eIF4E, *(R2)* ∼45-60 of 4E-BP2 with ∼135-150 of eIF4E, *(R3)* ∼78-82 of 4E-BP2 with ∼83-90 of eIF4E, *(R4)* ∼83-90 of 4E-BP2 with ∼76-83 of eIF4E, and *(R5)* ∼60-73 of 4E-BP2 with ∼60-80 of eIF4E (**Fig. 6b**). These contacts correspond to the following interactions: *R1* – residues adjacent of the 4E-BP2 canonical binding site with the root of the eIF4E *N*-IDR, *R2* – residues including the 4E-BP2 canonical binding site ^54^YXXXXLΦ^60^ and the turn between α2 helix and β5 sheet in eIF4E, *R3* – the 4E-BP2 secondary binding site ^78^IPGVT^82^ with the loop between α1 helix and β3 sheet in eIF4E, *R4* – residues adjacent to the 4E-BP2 secondary binding site and part of eIF4E α1 helix, and *R5* – the 4E-BP2 residues between the two binding sites and the α1 helix in eIF4E. These contacts are visualized in the color-coded 4E-BP2:eIF4E structures in **Fig. 6h**.

In the optimized ensemble the initial intermolecular contacts become more delocalized and are more dispersed around the contact regions (**Fig. 6c**). All existing contacts become enriched except for *R3*, the secondary binding site, consistent with the decrease in the fraction of conformers with a fixed secondary binding site (**Fig. 5d**). Two new regions of intermolecular contact emerge: *(RN1*) contacts of the *N*-IDR of eIF4E (residues 1-40) with the *N*-terminus of 4E-BP2 (residues 1-20), and *(RN1*) and sparse contacts between the *C*-terminus of 4E-BP2 (residues 110-120) and eIF4E, in particular residues 75-85 (**Fig. 6c & 6i**).

These enriched contacts are consistent with the binding-induced changes at the C-terminus of 4E-BP2 measured by time-resolved fluorescence spectroscopy, such as slower segmental dynamics and increased quenching dynamics (Smyth, Zhang et al. 2022). Binding induces changes to the NMR intensity ratios, which decrease ∼50-60% at residues ∼95-120, indicating that there are possible transient interactions with eIF4E (Lukhele, Bah et al. 2013). Electrostatics could explain the dynamic interaction mode of the *C*-terminus of 4E-BP2. This region has a predominantly negative net charge per residue (NCPR) while eIF4E has regions of positive NCPR at residues ∼40-65 and ∼105-120 of eIF4E (**Fig. S11**).

Deletion studies have shown that removing residues M1-N25 of eIF4E has no effect on the binding affinity of eIF4E to 4E-BP2, but deletion of residues M1-H33 increases the affinity by about two orders of magnitude (Abiko, Tomoo et al. 2007). Additionally, an NMR study of the interaction showed significant chemical shift changes starting at N25 of the eIF4E N-IDR when 4E-BP2 was bound (Lukhele, Bah et al. 2013). This suggests that residues N25-H33 form contacts with other parts of eIF4E that disrupt the binding of 4E-BP2. Interestingly, in the optimized ensemble the number of contacts between the eIF4E N-IDR and 4E-BP2 *N*-terminal region increases, but this apparent contradiction could be caused by allosteric effects. A possible scenario is that these intermolecular interactions with the eIF4E starting at residues N25 transmit allosteric changes to 4E-BP2 that make it less binding compatible. Furthermore, contacts between the canonical binding site ^54^YXXXXLΦ^60^ and eIF4E residues N25-H33 increase in the optimized ensemble, and these interactions would compete with those of the canonical binding site on eIF4E and weaken the interaction.

Clustering the 4E-BP2:eIF4E complex resulted in four clusters (**Fig. S10**) at the cutoff criterion (**Fig. S8b**). Intermolecular inter-residue contact maps of the clusters (**Fig. 6d-g**) paint a heterogeneous picture of the conformations comprising the ensemble. *Cluster 2* has the greatest number of *RN1* contacts but the second lowest number of *RN2* contacts amongst all the clusters (**Fig. 6d**). At the opposite end, *Cluster 4* has the greatest number of *RN2* contacts but the second fewest *RN1* contacts (**Fig. 6G**). Significant heterogeneity also exists amongst the four clusters regarding abundance of contacts in the initial regions *R1-R5* (**Table S4**). This heterogeneity reinforces the dynamic, heterogeneous nature of the complex which contributes to the fine-tuned binding affinity and its likely effects on post-translational modification in the bound state (Böhm, Imseng et al. 2021).

## DISCUSSION & CONCLUSIONS

In contrast to folded proteins, the sequences of IDPs encode for a relatively flat free-energy landscape characterized by a shallower, multi-funnel surface. Although there is a general understanding of how the amino acid sequence determines the populations of conformational ensembles of IDPs in their isolated state, less is known about how these ensembles are altered when interacting with folded proteins to form, in most cases, highly dynamic complexes. Especially for IDPs involved in signaling and regulatory processes, delineating these dynamic complexes is key for understanding their biological function. Here, we report one of the first conformational ensembles of an IDP in complex with its folded target, which was built using IDPConformerGenerator starting from a partial crystal structure and restrained by new experimental data. As restraints, we used single-molecule fluorescence measurements reporting on specific internal distances and on the global size of the complex, as well as SAXS and chemical shifts to drive secondary structure propensities.

smFRET provided restraints on chain dimensions and dynamics of 4E-BP2 at eight unique residue separations, in both the apo and the eIF4E-bound states. Across the entire chain, the free 4E-BP2 was either more compact or identical to a statistical coil prior, implying the existence of significant intramolecular contacts. In comparison to the previous apo 4E-BP2 ensemble (Tsangaris, Smyth et al. 2023), the new ensemble identified additional regions involved in compaction of sites separated by ∼70 residues. These additional intrachain contacts are most likely driven by electrostatic attractions, consistent with the conclusion from our previous study. Distance-based hierarchical clustering identified conformers with expansion surrounding the T37 and T46 phospho-sites. As these are the first two sites to be phosphorylated in hierarchical order, this local expansion could facilitate the access of regulatory kinases.

Intra-molecular FRET efficiency decreased upon binding for nearly all 4E-BP2 segments, and the magnitude of this effect is correlated with the fractional overlap of the respective segment with the known ‘binding region’ of 4E-BP2 (residues 54-82). This suggests that the expansion of the 4E-BP2 chain facilitates interaction of its canonical and secondary binding motifs with the two eIF4E binding sites, while it may also drive additional interacting regions. Notably, compared to the apo state, binding resulted in the compaction of segments **A** and **C** which probed the N-terminal and C-terminal parts of the chain respectively.

Due to the reasonably low computational expense of IDPConformerGenerator, nearly 100,000 conformations of the eIF4E:4E-BP2 complex were generated starting from a partial crystal structure (Peter, Igreja et al. 2015) to ensure a highly diverse initial pool. To account for the dynamic nature of the bipartite binding interface, some structural restraints were lifted and conformations were built by fixing only either one or both binding sites and then sampling the rest of the 4E-BP2 chain and the disordered regions of eIF4E using IDPConformerGenerator (Liu, Teixeira et al. 2023). A sub-ensemble of 100 of these structures was obtained by imposing agreement with the experimental fluorescence data using a new version of the Bayesian optimization method X-EISD (Lincoff, Haghighatlari et al. 2020). Since X-EISDv2 only compares experimental and back-calculated data, it is not limited to single-chain protein models; limitations only arise if the back-calculators do not accept multi-chain protein complexes. Furthermore, X-EISDv2 was designed with large-scale datasets in mind, so multiprocessing is built into the method to ensure fast scoring and optimization of the ∼100,000 all-atom conformer initial pool.

The refined ensemble of this IDP complex has a different fractional composition of *Canonical* only, *Secondary* only, and *Both* generation cases than the uniform distribution present in the initial ensemble. Namely, *Both* (59%) and *Canonical* (39%) become the dominant fractions at the expense of *Secondary* (2%). This indicates that the secondary binding site of the 4E-BP2 to eIF4E acts primarily in concert with the canonical binding site and that instances where only the secondary binding site is occupied are rare. Critically, it demonstrates that the crystal structure of the analogous 4E-BP1:eIF4E complex presentation of *Both* at 100% is not representative of the ensemble present in solution, providing a clear basis for the accessibility of regulatory kinases within the context of the 3 nM complex. It also aligns with deletion studies that report that the canonical but not the secondary binding site can bind to eIF4E in isolation (Paku, Umenaga et al. 2012). The resulting ensemble also shows increased contacts between residues in the *N*-IDR of eIF4E known to affect the binding of 4E-BP2 and the canonical binding surface of 4E-BP2. This is an interesting result as these contacts were not dominant in the initial pool but emerged as significant when restraints were imposed. Due to the generalizability of the IDPConformerGenerator and X-EISDv2 workflow, we can foresee the template method for generating dynamic complexes being feasible for other protein systems. Overall, this study exemplifies the mutually beneficial relationship between computations and experiments for defining disordered protein complexes, with advances in both likely to deepen our functional understanding of these complex heterogeneous molecular machines.

## MATERIALS and METHODS

### Sample preparation

Detailed protein expression, purification, and labelling procedures can be found in our previous publications(Dawson, Bah et al. 2020, Smyth, Zhang et al. 2022). Briefly, plasmids encoding for the SUMO fusion constructs of the 4E-BP2 or eIF4E proteins were expressed in BL21-codonplus (DE3) RIPL competent *Escherichia coli* cells (Agilent Technologies). Cells were lysed by sonication, protein was separated by crude lysate using a Ni-Sepharose column, followed by cleavage of the SUMO fusion and an additional Ni Sepharose separation. If purity was not sufficient the sample was further purified by FPLC, then was concentrated to 100 µM (glycerol added to 20% for eIF4E), flash frozen in liquid nitrogen and stored at -80°C.

Single-cysteine mutants of eIF4E and 4E-BP2 were labelled with Cy3 (Cytiva, PA23031) and Cy5 (Cytiva, Cat. no. PA25031), respectively, at a dye:protein molar ratio of 4:1 Double-cysteine mutants of 4E-BP2 were labelled simultaneously with AlexaFluor 488 (Invitrogen, Cat. no. A10254) and AlexaFluor 647 (Invitrogen, Cat. no. A20347) at a donor:acceptor:protein molar ratio of 3:2:1. Excess dye was separated from protein using size-exclusion chromatography on G-50 Sephadex resin (Sigma-Aldrich, Cat. no. G5080-10G). Fractions containing the labelled protein were pooled and concentrated to 100 µM (glycerol added to 20% for eIF4E), flash frozen in liquid nitrogen and stored at -80°C.

For burst smFRET measurements, the 4E-BP2 protein samples were diluted to concentrations of ∼50 pM in a PBS buffer containing 143 mM β-mercaptoethanol and 10 mM cysteamine hydrochloride to act as triplet state quenchers. TWEEN20 (Sigma-Aldrich, Cat. no. P2287) was also added at 0.005% (v/v) to prevent non-specific protein adsorption to the coverslip, the pH was adjusted to 7.4. For measurements of 4E-BP2 in complex with eIF4E, unlabeled eIF4E was added to the solution above to a final concentration of 2 µM. In a typical smFRET experiment, 30 µL of solution was dropped on the surface of a plasma-cleaned coverslip (Smyth, Zhang et al. 2022). All experiments were performed at 20°C.

Flow chambers enabling surface immobilization for TIRF smFRET measurements were constructed as described previously(Jain, Liu et al. 2012, Shivnaraine, Fernandes et al. 2016) (see SI for more details). The chambers were coated with bovine serum albumin (BSA) (Roy, Hohng et al. 2008), which for our protein system yielded very low non-specific adsorption due to the negative net charges of BSA (*Z* = −17), eIF4E (*Z* = −5), and 4E-BP2 (*Z* = −3). The empty chambers were rinsed by 0.4 mL of PBS buffer (137 mM NaCl, 2.3 mM KCl, 10 mM Na_2_HPO_4_, 1.8 mM KH_2_PO_4_, pH 7.4) followed by incubation with 50 µL of a 1 mg/mL mixture of 500:1 BSA:BSA-biotin for 20 minutes. After a PBS rinsing step, the chamber was incubated with 50 µL of 5 µg/mL streptavidin (MiliporeSigma, Cat. no. SA101) in PBS for 20 minutes. Another rinsing step was followed by incubation with 50 µL of 5 µg/mL of biotinylated Anti-FLAG antibody (Abcam, Cat. no. ab205606) in PBS for 20 minutes. The eIF4E:4E-BP2 complex was formed by mixing 5 nM of Cy3-labelled eIF4E-FLAG with 50 nM of Cy5-labelled 4E-BP2 in PBS; 50 µL of this mixture was incubated in the chamber for 5 minutes, preceded and followed by standard rinsing steps. Before TIRF imaging, 50 µL of fresh GODCAT oxygen scavenging solution (1 mg/mL glucose oxidase, 0.4 mg/mL catalase, 2 mM Trolox, 2 mM cyclooctatetraene (COT), pH 7.4) was flowed into the chamber to increase the molecular brightness and photostability of the Cy3 and Cy5 fluorophores (Aitken, Marshall et al. 2008).

### smFRET measurements

Intra-molecular smFRET experiments were performed on a custom-built multiparameter microscope described previously (Gomes, Krzeminski et al. 2020, Smyth, Zhang et al. 2022). The laser source was changed to a SuperK EXTREME EXW-12 PP (NKT Photonics, Birkød, Denmark) operated at 78.1 MHz, which outputs a broad spectrum of visible and infrared light (∼400-2400 nm) with a total output power of ∼4.6 W. In the visible 450-700 nm range, the SuperK laser outputs powers of 3-5 mW/nm with a master seed laser pulse of 5 ps. Alternating Laser Excitation (ALEX) was performed by synchronous modulation of the laser spectrum via an Acusto-Optical Tunable Filter (AOTF) (97-02885-04, Gooch and Housego, USA) which resulted in alternating excitation periods of 50 µs for the donor and the acceptor fluorophores. As such, FRET samples were excited at 488 nm and 633 nm and intensities of 23 kW/cm^2^ and 12 kW/cm^2^, respectively.

smFRET burst data was analyzed as described previously (Gomes, Krzeminski et al. 2020, Smyth, Zhang et al. 2022). Thousands of bursts for each sample were sorted into donor-only, acceptor-only, and dual-labeled populations. For each dual-labelled burst, the FRET efficiency was calculated as:

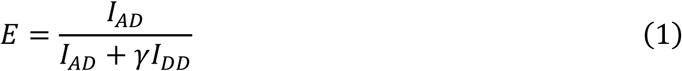

where *I*_*dd*_ and *I*_*Ad*_ are numbers of photons detected in the donor and the acceptor channels, respectively, following excitation of the donor, and *γ* is a correction factor for different detection efficiencies and fluorescence quantum yields of the two species. *I*_*dd*_ and *I*_*Ad*_ were corrected for background, spectral crosstalk, and direct excitation of the acceptor dye. Background was measured from a sample buffer containing everything except the FRET molecules.

Inter-molecular smFRET experiments were performed on a TIRF microscope (Liu, Mazouchi et al. 2010). Flow chambers containing immobilized protein complexes were imaged for a total of 100-1000 frames at a frame rate of 2-10 s^-1^. Dual-color images were captured by two cooled EMCCD cameras (DU-897BV and DU-10669, Andor, USA) from a sample area of 42 × 42 µm^2^. Donor excitation at 532 nm (*I* = 220 *W*/*cm*^2^) was used, except for the first and the last 10 frames when the acceptor was directly excitated at 633 nm (*I* = 38 *W*/*cm*^2^).

Dual-color TIRF image stacks were analyzed using a custom-written MATLAB analysis program (Zhou, Septien-Gonzalez et al. 2024), which detects the (*x,y*) coordinates of single emitters/particles in each channel and extracts their background-corrected intensity-time traces. Particles were classified as FRET particles if *i*) they were detected in each channel (e.g., *I_AA_*, *I_AD_* and *I_DD_*), and *ii*) their center-to-center distances in each channel did not exceed 1 pixel. FRET particles were further filtered by keeping only those where the acceptor or the donor dye photobleached before the end of the acquisition.

FRET efficiency was calculated using equation (*1*) for each TIRF frame using donor excitation in which both donor and acceptor fluorophores were active. The cumulative FRET histogram for all active frames of all FRET particles was fitted to a Gaussian distribution or, for high FRET values, to a non-normalized Beta distribution:

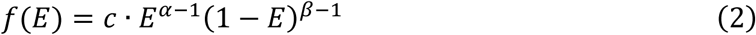

resulting in the mean FRET efficiency value 〈*E*〉 = α/α + β.

### IDP conformational ensembles: generation, refinement and analysis

The apo 4E-BP2 conformers were generated using IDPConformerGenerator, a software suite developed in the Forman-Kay lab (Teixeira, Liu et al. 2022). A total of 80,000 4E-BP2 conformations were generated with the default FASPR sidechain packing method (Huang, Pearce et al. 2020). Sidechains were protonated by using PDBFixer within the OpenMM suite (Eastman, Galvelis et al. 2024), then each conformer underwent a quick energy minimization simulation through OpenMM to resolve steric clashes. The statistical coil ensemble of 4E-BP2 was generated using the TraDES (Feldman and Hogue 2002), which uses a Lennard-Jones like potential to avoid steric clashes. 5,000 structures were generated using only coil-type dihedral angle sampling (Tsangaris, Smyth et al. 2023).

A conformational ensemble of the eIF4E:4E-BP2 complex was generated based on the 4E-BP1:eIF4E complex crystal structure (PDB ID: 4UED)(Peter, Igreja et al. 2015), which contains coordinates for residues 50-83 of 4E-BP2 and all residues of eIF4E except 1-32 and 205-211. The eIF4E:4E-BP2 complex ensemble was generated based on the crystal structure where three individual cases were modelled: **1**) only the canonical binding site of 4E-BP2 was fixed (residues 54-60), **2**) only the secondary binding site of 4E-BP2 was fixed (78-82), and **3**) both binding sites fixed (residues 54-60 and 78-82). The remaining residues of 4E-BP2 and the unresolved regions of eIF4E were sampled with the LDRS module (Liu, Teixeira et al. 2023) of the IDPConformerGenerator suite. For each case, 33,000 conformers were generated, with the final ensemble containing 99,000 conformers.

The original X-EISD method is a Bayesian approach that maximizes the log-likelihood of a disordered protein ensemble while accounting for the uncertainties in the back-calculation and experimental data. It is formulated as a general Bayesian model (Lincoff, Haghighatlari et al. 2020):

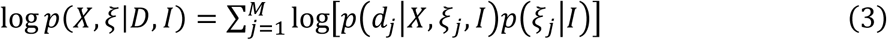

where log *p*(*X*, ζ|*D*, *I*) is the log-likelihood that an ensemble *X* composed of *N* conformers agrees with a set of *M* experimental values 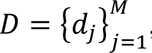, given back-calculation and experimental errors ζ, and prior information *I*. The new X-EISDv2 uses the same core logic as the original X-EISD where a user-defined sub-ensemble pool size, from which conformers are exchanged one at a time with replacement, the conformer swap is accepted if it leads to a higher total log-likelihood. X-EISDv2 has been re-written to be generalized to all protein systems, and the command-line interface has been designed to work as a standalone program that integrates into the IDPConformerGenerator and SPyCi-PDB workflow as a final scoring and reweighting protocol. Since X-EISDv2 is a scoring protocol at its core, the only requirements are that the back-calculated data format matches the experimental data, so it is not restricted to the type of biological system (i.e. multi-protein complexes). X-EISDv2 version 0.3.0 (github.com/THGLab/X-EISDv2/releases/tag/v0.3.) used in this study includes the normalization of experimental data so the number of experimental data points does not bias the final X-EISD score. Furthermore, custom-weighting based on the confidence of experimental data has been implemented, where a numerical multiplier (with a total sum of 1 for all multipliers) is used to allow bias towards more confident experimental data in the final X-EISD score value.

The mean FRET efficiency (〈*E*〉) for each conformer was calculated using accessible volume simulations(Sindbert, Kalinin et al. 2011, Kalinin, Peulen et al. 2012) by means of the *AvTraj*(Dimura, Peulen et al. 2016) and *MDTraj*(McGibbon, Beauchamp et al. 2015) Python packages; see Tsangaris et al. for more details (Tsangaris, Smyth et al. 2023). The hydrodynamic radius (*R_h_*) for each conformer was calculated using the *HullRad* Python program with default parameters(Fleming and Fleming 2018).

Contact maps were constructed by calculating the fraction of conformers at each inter-residue pair using a *C*_α_–*C*_α_ distance cutoff of 8 Å (*f*_*ij*_):

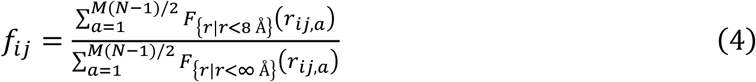

where *F*_*A*_(*a*) is an indicator function for the set *A* which outputs 1 if *a* ∈ *A* and 0 otherwise (Tsangaris, Smyth et al. 2023). For inter-molecular contacts between two proteins, *M* and *N* are the number of residues of each protein; for intramolecular contacts *M* = *N*. Difference contact maps were constructed by subtracting the fractions of the ensemble of interest *f*_*ij*,1_ from the reference ensemble *f*_*ij*,2_ at each residue separation.

## ACKNOWLEDGEMENTS

This work has been supported by the Natural Sciences and Engineering Research Council of Canada (NSERC) RGPIN-2023-04864 to C.C.G.; by the Canadian Institutes of Health Research #PJT-190060, NSERC RGPIN-2024-05725, and the Canada Research Chairs Program to J.D.F.-K.; and by the National Institutes of Health R01GM127627 to T.H.-G. and J.D.F.-K.

## REFERENCES

Abiko, F., K. Tomoo, A. Mizuno, S. Morino, H. Imataka and T. Ishida (2007). “Binding preference of eIF4E for 4E-binding protein isoform and function of eIF4E N-terminal flexible region for interaction, studied by SPR analysis.” Biochemical and biophysical research communications 355(3): 667–672.

Aitken, C. E., R. A. Marshall and J. D. Puglisi (2008). “An Oxygen Scavenging System for Improvement of Dye Stability in Single-Molecule Fluorescence Experiments.” Biophysical Journal 94(5): 1826–1835.

Alderson, T. R., I. Pritišanac, Đ. Kolarić, A. M. Moses and J. D. Forman-Kay (2023). “Systematic identification of conditionally folded intrinsically disordered regions by AlphaFold2.” Proceedings of the National Academy of Sciences 120(44): e2304302120.

Bah, A., R. M. Vernon, Z. Siddiqui, M. Krzeminski, R. Muhandiram, C. Zhao, N. Sonenberg, L. E. Kay and J. D. Forman-Kay (2015). “Folding of an intrinsically disordered protein by phosphorylation as a regulatory switch.” Nature 519(7541): 106–109.

Banko, J. L., M. Merhav, E. Stern, N. Sonenberg, K. Rosenblum and E. Klann (2007). “Behavioral alterations in mice lacking the translation repressor 4E-BP2.” Neurobiology of Learning and Memory 87(2): 248–256.

Baul, U., D. Chakraborty, M. L. Mugnai, J. E. Straub and D. Thirumalai (2019). “Sequence Effects on Size, Shape, and Structural Heterogeneity in Intrinsically Disordered Proteins.” The Journal of Physical Chemistry B 123(16): 3462–3474.

Böhm, R., S. Imseng, R. P. Jakob, M. N. Hall, T. Maier and S. Hiller (2021). “The dynamic mechanism of 4E-BP1 recognition and phosphorylation by mTORC1.” Molecular cell 81(11): 2403–2416. e2405.

Borgia, A., M. B. Borgia, K. Bugge, V. M. Kissling, P. O. Heidarsson, C. B. Fernandes, A. Sottini, A. Soranno, K. J. Buholzer and D. Nettels (2018). “Extreme disorder in an ultrahigh-affinity protein complex.” Nature 555(7694): 61–66.

Bottaro, S., T. Bengtsen and K. Lindorff-Larsen (2020). “Integrating molecular simulation and experimental data: a Bayesian/maximum entropy reweighting approach.” Structural bioinformatics: methods and protocols: 219–240.

Chen, J. and R. W. Kriwacki (2018). “Intrinsically disordered proteins: structure, function and therapeutics.” Journal of molecular biology 430(16): 2275.

Chowdhury, A., D. Nettels and B. Schuler (2023). “Interaction Dynamics of Intrinsically Disordered Proteins from Single-Molecule Spectroscopy.” Annual Review of Biophysics 52: 433–462.

Csizmok, V., A. V. Follis, R. W. Kriwacki and J. D. Forman-Kay (2016). “Dynamic protein interaction networks and new structural paradigms in signaling.” Chemical reviews 116(11): 6424–6462.

Dawson, J. E., A. Bah, Z. Zhang, R. M. Vernon, H. Lin, P. A. Chong, M. Vanama, N. Sonenberg, C. C. Gradinaru and J. D. Forman-Kay (2020). “Non-cooperative 4E-BP2 folding with exchange between eIF4E-binding and binding-incompatible states tunes cap-dependent translation inhibition.” Nat Commun 11(1): 3146.

Dimura, M., T. O. Peulen, C. A. Hanke, A. Prakash, H. Gohlke and C. A. Seidel (2016). “Quantitative FRET studies and integrative modeling unravel the structure and dynamics of biomolecular systems.” Current opinion in structural biology 40: 163–185.

Dogan, J., S. Gianni and P. Jemth (2014). “The binding mechanisms of intrinsically disordered proteins.” Physical Chemistry Chemical Physics 16(14): 6323–6331.

Eastman, P., R. Galvelis, R. P. Peláez, C. R. A. Abreu, S. E. Farr, E. Gallicchio, A. Gorenko, M. M. Henry, F. Hu, J. Huang, A. Krämer, J. Michel, J. A. Mitchell, V. S. Pande, J. P. Rodrigues, J. Rodriguez-Guerra, A. C. Simmonett, S. Singh, J. Swails, P. Turner, Y. Wang, I. Zhang, J. D. Chodera, G. De Fabritiis and T. E. Markland (2024). “OpenMM 8: Molecular Dynamics Simulation with Machine Learning Potentials.” The Journal of Physical Chemistry B 128(1): 109–116.

Feldman, H. J. and C. W. Hogue (2002). “Probabilistic sampling of protein conformations: new hope for brute force?” Proteins: Structure, Function, and Bioinformatics 46(1): 8–23.

Ferrie, J. J. and E. J. Petersson (2020). “A Unified De Novo Approach for Predicting the Structures of Ordered and Disordered Proteins.” The Journal of Physical Chemistry B 124(27): 5538–5548.

Fleming, P. J. and K. G. Fleming (2018). “HullRad: Fast calculations of folded and disordered protein and nucleic acid hydrodynamic properties.” Biophysical journal 114(4): 856–869.

Fletcher, C. M., A. M. McGuire, A.-C. Gingras, H. Li, H. Matsuo, N. Sonenberg and G. Wagner (1998). “4E binding proteins inhibit the translation factor eIF4E without folded structure.” Biochemistry 37(1): 9–15.

Fuxreiter, M. (2019). “Fold or not to fold upon binding — does it really matter?” Current Opinion in Structural Biology 54: 19–25.

Ghafouri, H., T. Lazar, A. Del Conte, L. G. Tenorio Ku, P. Consortium, P. Tompa, S. C. E. Tosatto and A. M. Monzon (2023). “PED in 2024: improving the community deposition of structural ensembles for intrinsically disordered proteins.” Nucleic Acids Research 52(D1): D536–D544.

Gingras, A.-C., S. P. Gygi, B. Raught, R. D. Polakiewicz, R. T. Abraham, M. F. Hoekstra, R. Aebersold and N. Sonenberg (1999). “Regulation of 4E-BP1 phosphorylation: a novel two-step mechanism.” Genes & development 13(11): 1422–1437.

Gomes, G.-N. W., M. Krzeminski, A. Namini, E. W. Martin, T. Mittag, T. Head-Gordon, J. D. Forman-Kay and C. C. Gradinaru (2020). “Conformational ensembles of an intrinsically disordered protein consistent with NMR, SAXS, and single-molecule FRET.” Journal of the American Chemical Society 142(37): 15697–15710.

Gopich, I. V. and A. Szabo (2012). “Theory of the energy transfer efficiency and fluorescence lifetime distribution in single-molecule FRET.” Proceedings of the National Academy of Sciences 109(20): 7747–7752.

Hadži, S., R. Loris and J. Lah (2021). “The sequence–ensemble relationship in fuzzy protein complexes.” Proceedings of the National Academy of Sciences 118(37): e2020562118.

Holehouse, A. S. and B. B. Kragelund (2024). “The molecular basis for cellular function of intrinsically disordered protein regions.” Nature Reviews Molecular Cell Biology 25(3): 187–211.

Huang, X., R. Pearce and Y. Zhang (2020). “FASPR: an open-source tool for fast and accurate protein side-chain packing.” Bioinformatics 36(12): 3758–3765.

Iljina, M., G. A. Garcia, M. H. Horrocks, L. Tosatto, M. L. Choi, K. A. Ganzinger, A. Y. Abramov, S. Gandhi, N. W. Wood and N. Cremades (2016). “Kinetic model of the aggregation of alpha-synuclein provides insights into prion-like spreading.” Proceedings of the National Academy of Sciences 113(9): E1206–E1215.

Jain, A., R. Liu, Y. K. Xiang and T. Ha (2012). “Single-molecule pull-down for studying protein interactions.” Nature protocols 7(3): 445–452.

Kalinin, S., T. Peulen, S. Sindbert, P. J. Rothwell, S. Berger, T. Restle, R. S. Goody, H. Gohlke and C. A. Seidel (2012). “A toolkit and benchmark study for FRET-restrained high-precision structural modeling.” Nature methods 9(12): 1218–1225.

LeBlanc, S. J., P. Kulkarni and K. R. Weninger (2018). “Single molecule FRET: A powerful tool to study intrinsically disordered proteins.” Biomolecules 8(4): 140.

Lincoff, J., M. Haghighatlari, M. Krzeminski, J. M. Teixeira, G.-N. W. Gomes, C. C. Gradinaru, J. D. Forman-Kay and T. Head-Gordon (2020). “Extended experimental inferential structure determination method in determining the structural ensembles of disordered protein states.” Communications chemistry 3(1): 74.

Liu, B., A. Mazouchi and C. C. Gradinaru (2010). “Trapping Single Molecules in Liposomes: Surface Interactions and Freeze− Thaw Effects.” The Journal of Physical Chemistry B 114(46): 15191–15198.

Liu, Z. H., J. M. Teixeira, O. Zhang, T. E. Tsangaris, J. Li, C. C. Gradinaru, T. Head-Gordon and J. D. Forman-Kay (2023). “Local disordered region sampling (LDRS) for ensemble modeling of proteins with experimentally undetermined or low confidence prediction segments.” Bioinformatics 39(12): btad739.

Lukhele, S., A. Bah, H. Lin, N. Sonenberg and J. D. Forman-Kay (2013). “Interaction of the eukaryotic initiation factor 4E with 4E-BP2 at a dynamic bipartite interface.” Structure 21(12): 2186–2196.

McGibbon, R. T., K. A. Beauchamp, M. P. Harrigan, C. Klein, J. M. Swails, C. X. Hernández, C. R. Schwantes, L.-P. Wang, T. J. Lane and V. S. Pande (2015). “MDTraj: a modern open library for the analysis of molecular dynamics trajectories.” Biophysical journal 109(8): 1528–1532.

Metskas, L. A. and E. Rhoades (2020). “Single-molecule FRET of intrinsically disordered proteins.” Annual review of physical chemistry 71: 391–414.

Mollica, L., L. M. Bessa, X. Hanoulle, M. R. Jensen, M. Blackledge and R. Schneider (2016). “Binding mechanisms of intrinsically disordered proteins: theory, simulation, and experiment.” Frontiers in molecular biosciences 3: 52.

Naudi-Fabra, S., M. Tengo, M. R. Jensen, M. Blackledge and S. Milles (2021). “Quantitative description of intrinsically disordered proteins using single-molecule FRET, NMR, and SAXS.” Journal of the American Chemical Society 143(48): 20109–20121.

Nojima, H., C. Tokunaga, S. Eguchi, N. Oshiro, S. Hidayat, K.-i. Yoshino, K. Hara, N. Tanaka, J. Avruch and K. Yonezawa (2003). “The mammalian target of rapamycin (mTOR) partner, raptor, binds the mTOR substrates p70 S6 kinase and 4E-BP1 through their TOR signaling (TOS) motif.” Journal of Biological Chemistry 278(18): 15461–15464.

Omidi, A., M. H. Møller, N. Malhis, J. M. Bui and J. Gsponer (2024). “AlphaFold-Multimer accurately captures interactions and dynamics of intrinsically disordered protein regions.” Proceedings of the National Academy of Sciences 121(44): e2406407121.

Paku, K. S., Y. Umenaga, T. Usui, A. Fukuyo, A. Mizuno, Y. In, T. Ishida and K. Tomoo (2012). “A conserved motif within the flexible C-terminus of the translational regulator 4E-BP is required for tight binding to the mRNA cap-binding protein eIF4E.” Biochemical Journal 441(1): 237–245.

Peter, D., C. Igreja, R. Weber, L. Wohlbold, C. Weiler, L. Ebertsch, O. Weichenrieder and E. Izaurralde (2015). “Molecular architecture of 4E-BP translational inhibitors bound to eIF4E.” Molecular cell 57(6): 1074–1087.

Ravera, E., L. Sgheri, G. Parigi and C. Luchinat (2016). “A critical assessment of methods to recover information from averaged data.” Physical Chemistry Chemical Physics 18(8): 5686–5701.

Rouse Jr, P. E. (1953). “A theory of the linear viscoelastic properties of dilute solutions of coiling polymers.” The Journal of Chemical Physics 21(7): 1272–1280.

Roy, R., S. Hohng and T. Ha (2008). “A practical guide to single-molecule FRET.” Nature methods 5(6): 507–516.

Schalm, S. S., D. C. Fingar, D. M. Sabatini and J. Blenis (2003). “TOS motif-mediated raptor binding regulates 4E-BP1 multisite phosphorylation and function.” Current biology 13(10): 797–806.

Sharma, R., Z. Raduly, M. Miskei and M. Fuxreiter (2015). “Fuzzy complexes: Specific binding without complete folding.” FEBS Letters 589(Part A): 2533–2542.

Shivnaraine, R. V., D. D. Fernandes, H. Ji, Y. Li, B. Kelly, Z. Zhang, Y. R. Han, F. Huang, K. S. Sankar and D. N. Dubins (2016). “Single-molecule analysis of the supramolecular organization of the M2 muscarinic receptor and the Gαi1 protein.” Journal of the American Chemical Society 138(36): 11583–11598.

Sindbert, S., S. Kalinin, H. Nguyen, A. Kienzler, L. Clima, W. Bannwarth, B. Appel, S. Müller and C. A. Seidel (2011). “Accurate distance determination of nucleic acids via Forster resonance energy transfer: implications of dye linker length and rigidity.” Journal of the American Chemical Society 133(8): 2463–2480.

Smyth, S. (2024). Characterization of Disordered Protein States and Complexes using Single-molecule Fluorescence Spectroscopy PhD thesis, University of Toronto.

Smyth, S., Z. Zhang, A. Bah, T. E. Tsangaris, J. Dawson, J. D. Forman-Kay and C. C. Gradinaru (2022). “Multisite phosphorylation and binding alter conformational dynamics of the 4E-BP2 protein.” Biophys J 121(16): 3049–3060.

Sonenberg, N. and A. G. Hinnebusch (2009). “Regulation of translation initiation in eukaryotes: mechanisms and biological targets.” Cell 136(4): 731–745.

Tee, A. R. and C. G. Proud (2002). “Caspase cleavage of initiation factor 4E-binding protein 1 yields a dominant inhibitor of cap-dependent translation and reveals a novel regulatory motif.” Molecular and cellular biology 22(6): 1674–1683.

Teixeira, J. M., Z. H. Liu, A. Namini, J. Li, R. M. Vernon, M. Krzeminski, A. A. Shamandy, O. Zhang, M. Haghighatlari and L. Yu (2022). “IDPConformerGenerator: A flexible software suite for sampling the conformational space of disordered protein states.” The Journal of Physical Chemistry A 126(35): 5985–6003.

Tsang, B., I. Pritišanac, S. W. Scherer, A. M. Moses and J. D. Forman-Kay (2020). “Phase Separation as a Missing Mechanism for Interpretation of Disease Mutations.” Cell 183(7): 1742–1756.

Tsangaris, T. E., S. Smyth, G.-N. W. Gomes, Z. H. Liu, M. Milchberg, A. Bah, G. A. Wasney, J. D. Forman-Kay and C. C. Gradinaru (2023). “Delineating Structural Propensities of the 4E-BP2 Protein via Integrative Modeling and Clustering.” The Journal of Physical Chemistry B 127(34): 7472–7486.

Uversky, V. N. (2025). “How to drug a cloud? Targeting intrinsically disordered proteins.” Pharmacological Reviews 77(2).

Uversky, V. N., C. J. Oldfield and A. K. Dunker (2008). “Intrinsically Disordered Proteins in Human Diseases: Introducing the D2 Concept.” Annual Review of Biophysics 37(1): 215–246.

Wiggers, F., S. Wohl, A. Dubovetskyi, G. Rosenblum, W. Zheng and H. Hofmann (2021). “Diffusion of a disordered protein on its folded ligand.” Proceedings of the National Academy of Sciences 118(37): e2106690118.

Wright, P. E. and H. J. Dyson (2009). “Linking folding and binding.” Current Opinion in Structural Biology 19(1): 31–38.

Zhou, X., H. Septien-Gonzalez, S. Husaini, R. J. Ward, G. Milligan and C. C. Gradinaru (2024). “Diffusion and Oligomerization States of the Muscarinic M1 Receptor in Live Cells – The Impact of Ligands and Membrane Disruptors.” Manuscript submitted for publication.

