## Supplemental Information for "Conformational Ensembles of the Disordered 4E-BP2:eIF4E Complex Restrained by smFRET Experiments"

### SUPPLEMENTARY METHODS

#### 1. Sample Preparation

#### 1.1 Protein Expression and Purification

Detailed protein expression, purification, and labelling procedures can be found in our previous publications (Dawson, Bah et al. 2020, Smyth, Zhang et al. 2022). Briefly, BL21-codonplus (DE3) RIPL competent *Escherichia coli* cells (Agilent Technologies) were transformed with plasmids encoding for the SUMO fusion constructs of the 4E-BP2 or eIF4E proteins. Cells were cultured in 1 L of LB medium at 37ºC shaking at 240 rpm until an $OD_{600}$ ~0.6-0.8 was reached, induced with 1 mM IPTG, and expressed at 16ºC for ~16 hours. Cells expressing 4E-BP2 were lysed by sonication in a denaturing phosphate buffer (20 mM $NH_2PO_4$, 500 mM NaCl, 20 mM Imidazole, 6 M GdmCl (guanidinium chloride), and 5 mM β-met, pH 7.4). Cells expressing eIF4E were lysed in a non-denaturing phosphate buffer (20 mM $NH_2PO_4$, 500 mM NaCl, 20 mM Imidazole, 5 mM β-met, ~0.5 mg DNASE1, and cOmplete, EDTA-free protease inhibitor cocktail (Roche), pH 7.4).

Protein was separated from the crude lysate by binding to a Ni-Sepharose column for 30 minutes, the column was washed with at least 20 column volumes (CVs) of lysis buffer followed by elution with 10 or more CVs of elution buffer (lysis buffer with 400 mM Imidazole). The elution fraction was dialyzed against 4L of Tris buffer (50 mM Tris-HCl, 250 mM NaCl, 10 mM Imidazole, and 5 mM β-met) with ~100 µg ULP1 protease added to cleave the SUMO fusion tag. The SUMO tag was separated from the target protein by binding to a Ni Sepharose, protein concentration was measured at $A_{280}$ or with the Bradford assay. If the protein was not pure as assessed by SDS-PAGE, FPLC with a HiLoad Superdex 75 PG gel filtration column (GE Healthcare) was performed. The purity and identity of the proteins were verified by ESI-MS.

Proteins were buffer exchange into PBS buffer, concentrated to 100 µM (glycerol added to 20% for eIF4E) and flash frozen in liquid nitrogen and stored at -80ºC.

### 1.2 Fluorescence Labelling

Single-cysteine mutants of eIF4E (I35C and T205C) and 4E-BP2 S14C/C35S/C73S (C14) and C35S/C73S/C121 (C121) were labelled with Cy3 maleimide (Cytiva, PA23031) and Cy5 maleimide (Cytiva, Cat. no. PA25031) respectively at a dye:protein molar ratio of 4:1. Double-cysteine 4E-BP2 mutants (C0/H32C/C35S/C73S (C0C32/A), H32C/C35S/C73S/S91C (C32C91/B), C35S/C73S/S91C/C121 (C91C121/C), C0/C35S/C73S/S91C (C0C91/AB), H32C/C35S/C73S/C121 (C32C121/BC), C0/C35S/C73S/C121 (C0C121/ABC), S14C/C35S (C14C73/β-fold), C35S/C121 (C73C121/C-term)) were labelled simultaneously with Alexa Fluor 488 maleimide (Invitrogen, Cat. no. A10254) and Alexa Fluor 647 maleimide (Invitrogen, Cat. no. A20347) at a donor:acceptor:protein molar ratio of 3:2:1. Prior to labelling, the PBS buffer was exchanged four times into the labelling buffer (50 mM $NH_2PO_4$, 250 mM NaCl, 2 mM tris(2-carboxyethyl)phosphine hydrochloride (TCEP-HCl, Bio Basics, Cat. no. TB0974), pH 7.0) using a 3 kDa MWCO centrifugal filter (Milipore, Cat. no. UFC900308) to remove β-mercaptoethanol. Maleimide reactive dyes were dissolved to 20 mM in anhydrous dimethyl sulfoxide (DMSO) and stored at -20ºC for up to 6 months. After adding the maleimide reactive dyes the sample was flushed with Argon gas for one minute, sealed and reacted overnight at 4ºC with gently shaking. The unreacted dye was quenched by adding 10 mM dithiothreitol, excess dye was separated from protein using size-exclusion chromatography on G-50 Sephadex resin (Sigma-Aldrich, Cat. no. G5080-10G). Fractions containing the labelled protein were pooled and concentrated to 100 µM (glycerol added to 20% for eIF4E), and flash frozen in liquid nitrogen, and stored at -80ºC.

### 1.3 Flow Chambers for TIRF Measurements

Flow chambers enabling surface immobilization for TIRF smFRET measurements were constructed based on the protocols described previously(Jain, Liu et al. 2012, Shivnaraine, Fernandes et al. 2016). Glass cover-slides and No. 1 coverslips (VWR, Cat. no. 48366-089) were cleaned by 30 minutes of sonication in each of the following solutions: 1% Hellmanex (Sigma-Aldrich, Cat. no. Z805939-1EA) solution, Mili-Q water, 1M KOH (Fisher Scientific, Cat. no. 1310-58-3), acetonitrile (Sigma-Aldrich, Cat. no. 34851), 1M KOH, and MeOH (Sigma-Aldrich, Cat. no. 34860). The cover-slides and coverslips were blown dry with Argon before being joined together by placing a parafilm (MiliporeSigma, Cat. no. P7793) cutout between the cover-slide and coverslip. They were then heated on a hotplate for ~30 seconds until the parafilm melted, fusing the components together.

### 2. smFRET Corrections and Data Analysis

Leakage ($Lk$) and direct-excitation ($Dir$) corrections were obtained from measurements of 50 nM Cy3-ssDNA and 50 nM Cy5-ssDNA, respectively. Corrected FRET efficiencies were calculated by subtracting $F_{Lk} = Lk \cdot I_D$ and $F_{Dir} = Dir \cdot I_A$ from the numerator and denominator of equation (*1*), respectively. The correction factor $\gamma$ in equation (*1*) accounts for the difference in the apparent brightness (detection efficiency and fluorescence quantum yield) between the two donor and acceptor dyes. To estimate $\gamma$, two dsDNA constructs with lengths of 17 and 13 base pairs (bp) were used. Each are end-labelled with Cy 3 and Cy5 and have FRET efficiencies of ~30% and ~50%, respectively. The gamma factor was determined by using a procedure described previously (Lee, Kapanidis et al. 2005). The shot-noise limit ($\sigma_{sn}$) for each burst-based smFRET histogram provided the histogram width expected if shot-noise was the only contribution to broadening. It was computed for each smFRET histogram using the equation:

$$\sigma_{sn} = \sqrt{\frac{\langle E\rangle(1-\langle E\rangle)}{\langle N\rangle}} \qquad (S1)$$

Where $\langle E\rangle$ is the average FRET efficiency of the smFRET histogram obtained by Gaussian fitting and $\langle N\rangle$ is the average number of photons per burst.

### 3. Ensemble Calculations of Hydrodynamic Radius

*HullRad* uses a convex hull model to estimate the hydrodynamic volume of molecules and has a significantly reduced computation cost relative to other commonly used methods, such as Hydropro (Ortega, Amorós et al. 2011),. It also works well for folded, flexible, and disordered proteins (Fleming and Fleming 2018). The back-calculated $R_h$ values have an uncertainty of +/-5% given that *HullRad* has a reported 5% error in predicting $R_h$. The experimental uncertainty was estimated from several measurements Rhodamine 6G, a dye used for calibration of the confocal volume geometric parameters. The ratio of the standard deviation to the mean of the diffusion coefficients extracted from fitting the Rhodamine 6G data was 5%. Thus, the experimental $R_h$ value was assigned an uncertainty of +/-1.7 Å, based on the experimental value of 34.5 Å.

### SUPPLEMENTARY FIGURES

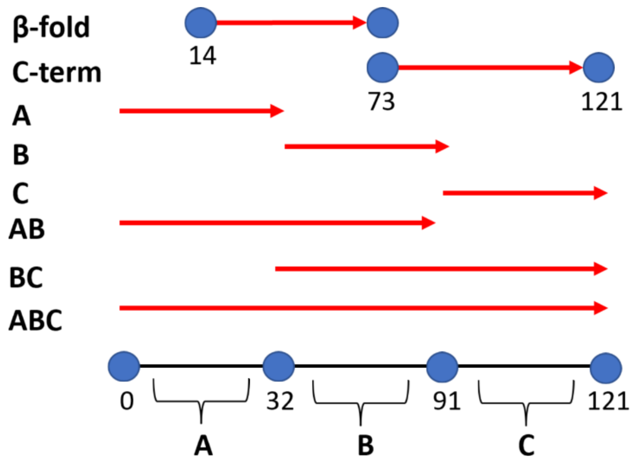


**Figure S1.** Schematic representation of the 4E-BP2 intramolecular smFRET constructs. The sequence was divided into three segments based on locations of cysteine mutations at positions 0, 32, 91, and 121. All nonredundant combinations of double-cysteine mutations were taken resulting in constructs ***A***, ***B***, ***C***, ***AB***, ***BC***, and ***ABC***. Two additional constructs, the ***β-fold*** (14-73) and ***C-term*** (73-121), were also used.

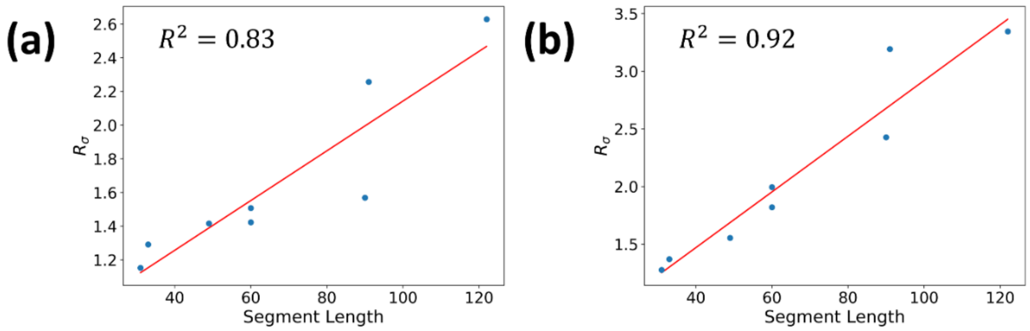


**Figure S2.** The correlation between the segment length (# of residues) probed by FRET in 4E-BP2 and the excess broadening of the experimental FRET distributions compared to the shot-noise limit ($R_\sigma = \sigma_{obs}/\sigma_{sn}$) for **(a)** apo 4E-BP2 and **(b)** 4E-BP2 bound to eIF4E.

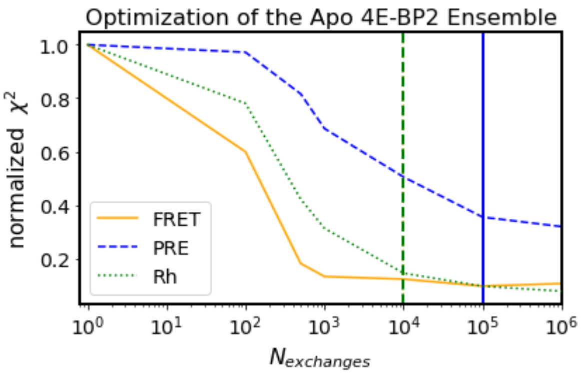


**Figure S3.** X-EISD optimization of the Apo 4E-BP2 ensemble. As the number of conformer exchanges ($N_{exch}$) increases, the normalized $\chi^2$ between back-calculated and experimental data decreases. The vertical lines indicate the optimum (knee-point) $N_{exch}$ values for FRET (yellow), $R_h$ (green) and PRE (blue). This optimization was done with a FRET weighting factor of $\Omega = 50$ (see Methods).

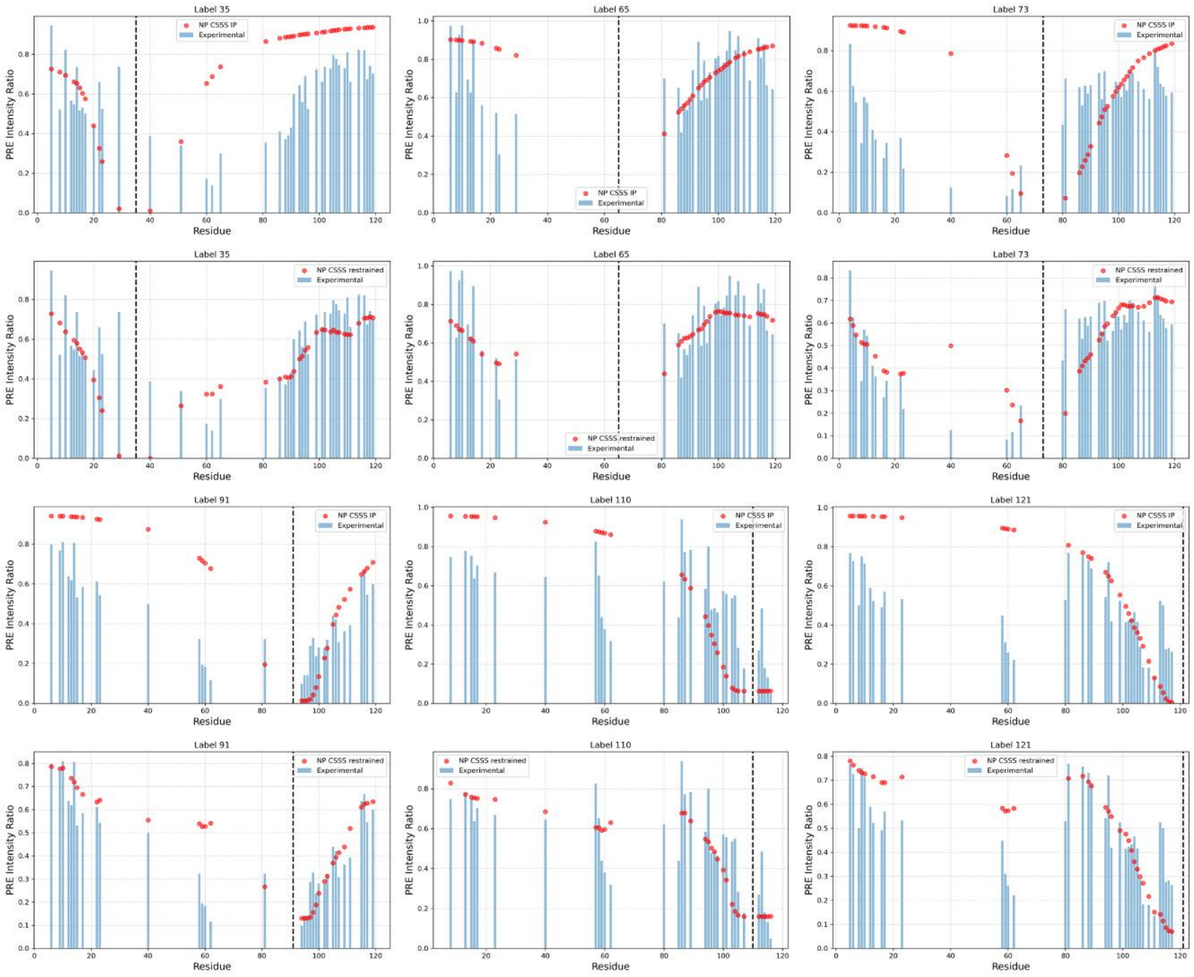


**Figure S4.** Comparison between the experimental PRE intensity ratios at 6 labeling positions in 4E-BP2, and those back calculated either from the initial pool (IP) of conformers, or from the X-EISD optimized ensemble (restrained).

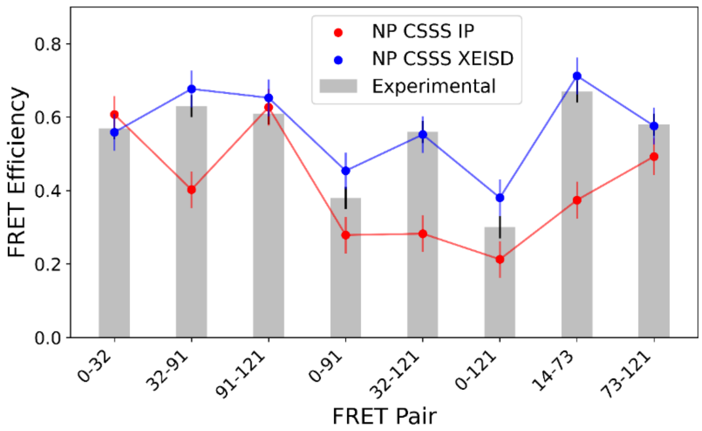


**Figure S5.** Comparison between the experimental FRET efficiencies used for restraining the non-phospho (NP) apo 4E-BP2 ensemble (grey bars), those back calculated from the initial pool (blue), and those back calculated from the X-EISD-optimized ensemble (red). Error bars are ±0.03 on the experimental and ±0.05 on the back calculated values. CSSS represents ensembles calculated by custom secondary structure sampling where the secondary structure propensity is biased by NMR chemical shifts.

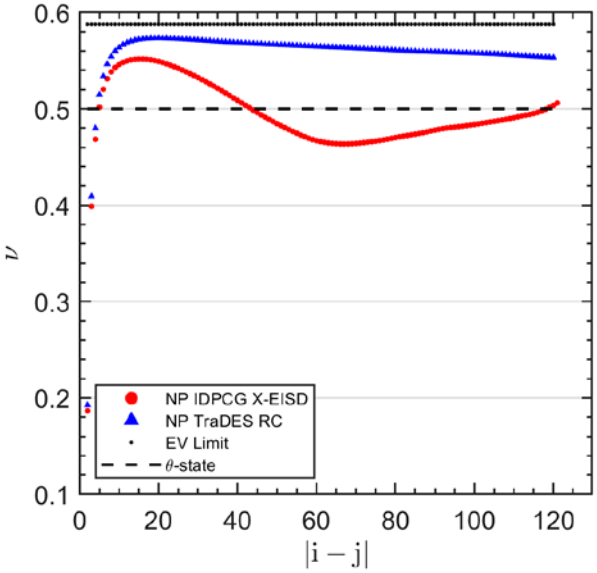


**Figure S6.** Per-residue scaling exponents of the optimized apo 4E-BP2 X-EISD ensemble (blue triangles), the TraDES RC ensemble (red dots), excluded volume (EV) limit (dots) and $\theta$-solvent polymers (dashed). The per-residue scaling exponent was calculated from $R_{|i-j|} = \sqrt{2l_p b}\,|i-j|^{\nu}$ by plotting $\nu(|i-j|) =$

$\log \frac{R_{|i-j|}}{\sqrt{2 l_p b}} / |i-j|$, where $b = 3.8$ Å is the distance between bonded $C_\alpha$ atoms and $l_p = 4$ Å is the persistence length.

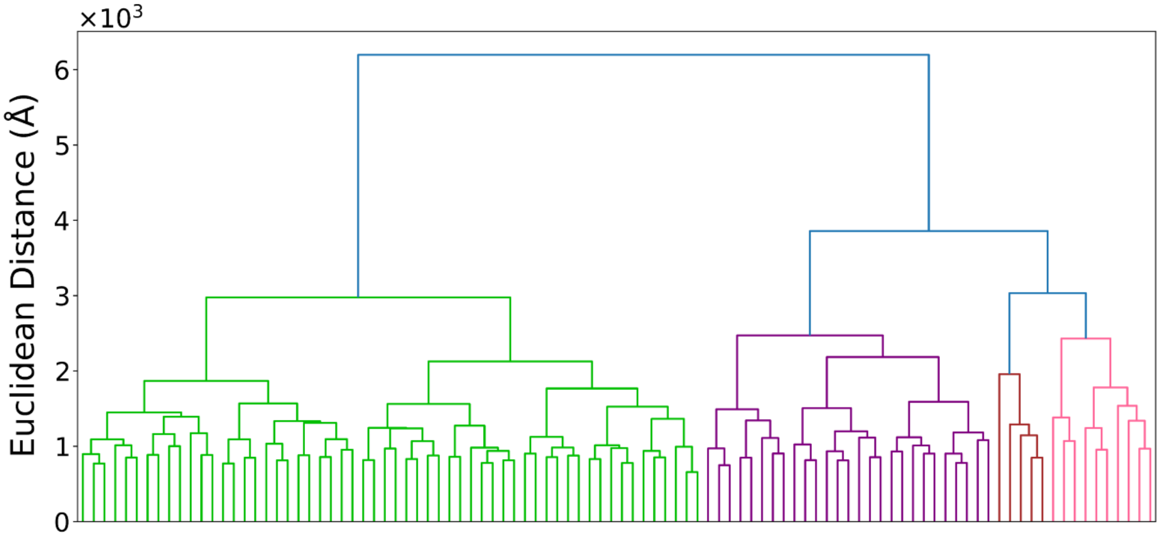


**Figure S7.** The dendrogram for agglomerative hierarchical clustering of the optimized apo 4E-BP2 complex ensemble; see (Tsangaris, Smyth et al. 2023) for more details. The resulting clusters are *Cluster 1* (green), *Cluster 2* (purple), *Cluster 3* (brown), and *Cluster 4* (pink).

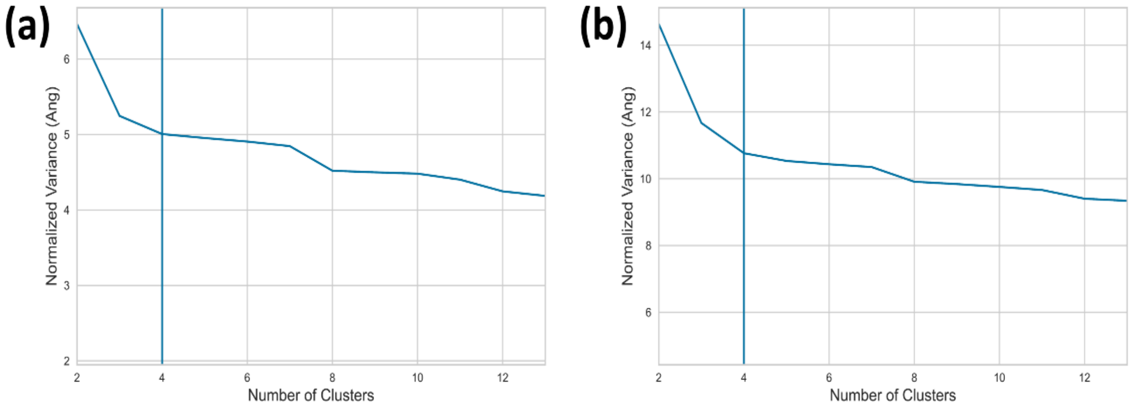


**Figure S8.** Normalized variance metric (Tsangaris, Smyth et al. 2023) used for determining the cutoff in determining the number of clusters for **(a)** optimized apo 4E-BP2 ensemble and **(b)** the optimized 4E-BP2: eIF4E complex ensemble. Note that for **(a)** *Clusters 3* and *4* were combined due to the low population (5%) of *Cluster 3*.

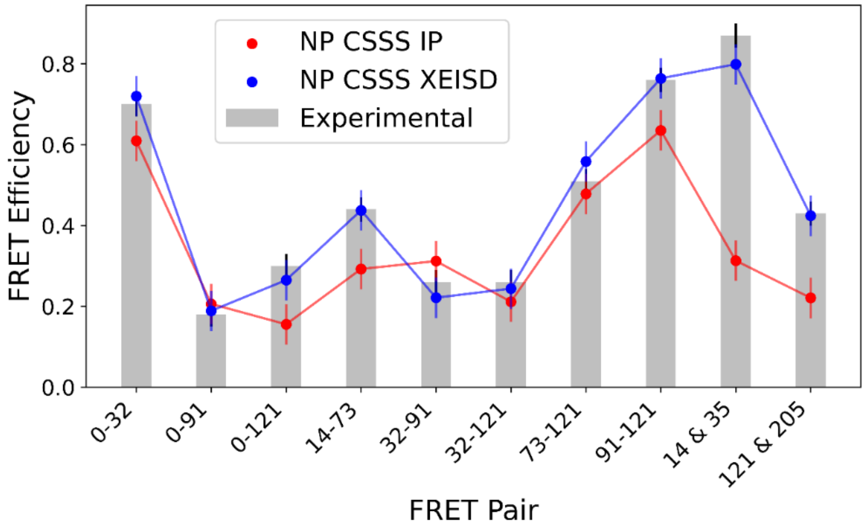


**Figure S9.** Comparison between the experimental FRET efficiencies used for restraining the conformational ensemble of the 4E-BP2:eIF4E complex (grey bars), those back calculated from the initial pool (red dots), and those back calculated from the optimized ensemble (blue dots). Error bars are ±0.03 on the experimental and ±0.05 on the back calculated values. CSSS represents ensembles calculated by custom secondary structure sampling where the secondary structure propensity is biased by NMR chemical shifts.

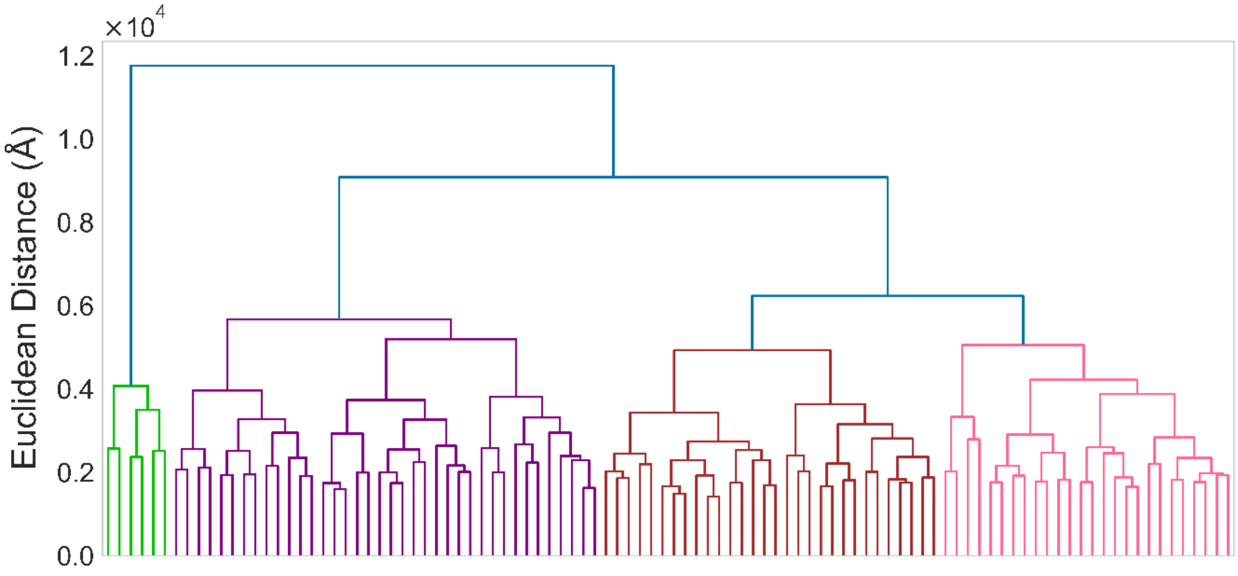

**Figure S10.** The dendrogram for agglomerative hierarchical clustering of the optimized 4E-BP2:eIF4E complex ensemble. The resulting clusters are *Cluster 1* (green), *Cluster 2* (purple), *Cluster 3* (brown), and *Cluster 4* (pink).

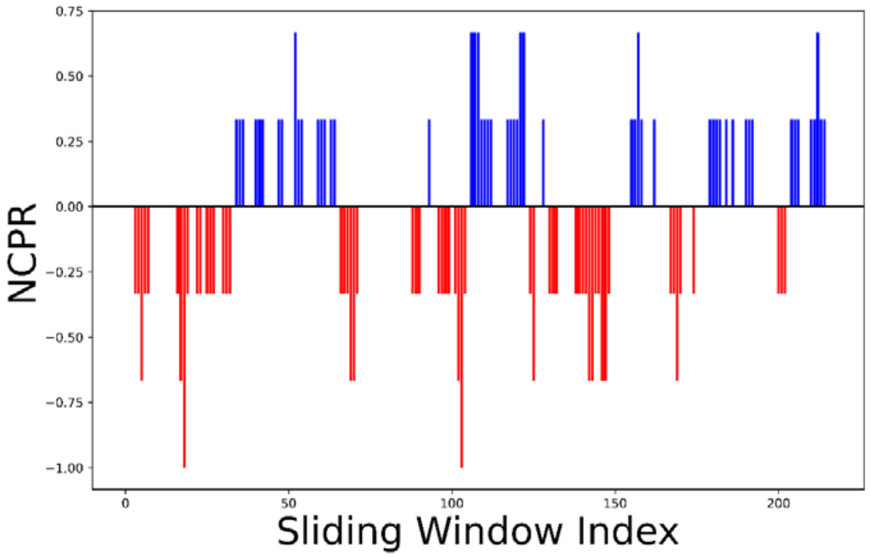


**Figure S11.** Net-charge-per-residue (NCPR) indexes for eIF4E calculated using a five-residue sliding-window (Holehouse, Das et al. 2017) across the amin-acid sequence of eIF4E. The charge of each amino acid was taken to be the most populated charge state at pH 7 using the pKa of each amino acid.

### SUPPLEMENTARY TABLES

**Table S1.** Mean FRET efficiencies, experimental ($\langle E \rangle_{exp}$)[a] and predicted ($\langle E \rangle_{RC}$)[b], for different 4E-BP2 sequence segments in the Apo and eIF4E-bound states

| | **4E-BP2** | | **4E-BP2:eIF4E** |
|---|---|---|---|
| | $\langle \boldsymbol{E} \rangle_{\boldsymbol{exp}}$ $(\pm 0.02)$ | $\langle \boldsymbol{E} \rangle_{\boldsymbol{RC}}$ $(\pm 0.03)$ | $\langle \boldsymbol{E} \rangle_{\boldsymbol{exp}}$ $(\pm 0.02)$ |
| **A**<br>(C0/H32C) | 0.57 | 0.61 | 0.70 |
| **B**<br>(H32C/S91C) | 0.63 | 0.43 | 0.26 |
| **C**<br>(S91C/C121) | 0.61 | 0.62 | 0.76 |
| **AB**<br>(C0/S91C) | 0.38 | 0.32 | 0.18 |
| **BC**<br>(H32C/C121) | 0.56 | 0.32 | 0.26 |
| **ABC**<br>(C0/C121) | 0.30 | 0.26 | 0.30 |
| **β-fold**<br>(S14C/C73) | 0.67 | 0.43 | 0.44 |
| **C-term**<br>(C73/C121) | 0.58 | 0.49 | 0.51 |

[a] The mean FRET efficiency obtained by Gaussian fitting of experimental smFRET histograms (**Fig. 2** and **Fig. 4** in the main text).
[b] The mean FRET efficiency predicted using the TraDES RC ensemble of Apo 4E-BP2 (see **Methods** in the main text).

**Table S2.** Comparison between mean FRET efficiencies of various 4E-BP2 segments predicted using a previous optimized ensemble[a] and experimental values measured in this study

| | $\langle E \rangle_{Tsangaris}$ ($\pm 0.03$) | $\langle E \rangle_{exp}$ ($\pm 0.02$) |
|---|---|---|
| **A** (C0/H32C) | 0.61 | 0.57 |
| **C** (S91C/C121) | 0.66 | 0.61 |
| **AB** (C0/S91C) | 0.37 | 0.38 |
| **BC** (H32C/C121)* | ***0.42*** | ***0.56*** |
| **ABC** (C0/C121) | 0.26 | 0.30 |
| **β-fold** (S14C/C73)* | ***0.40*** | ***0.67*** |

[a] For details about this optimized ensemble see (Tsangaris, Smyth et al. 2023).
*The prediction is not in agreement with the experimental mean FRET efficiency within uncertainties.

**Table S3.** Measured and calculated hydrodynamic radii ($R_h$, in Angstroms) of the Apo 4E-BP2 and the 4E-BP2:eIF4E complex[a]

| | **Apo 4E-BP2** | **4E-BP2:eIF4E complex** |
|---|---|---|
| **Experiment (FCS)** | $24.8 \pm 1.2$ | $34.5 \pm 1.4$ |
| **Initial Pool** | $30.5 \pm 1.5$ | $39.5 \pm 2.0$ |
| **Optimized Ensemble** | $26.4 \pm 1.3$ | $35.8 \pm 1.8$ |

[a] The experimental values were measured by FCS (fluorescence correlation spectroscopy). The back calculated values were obtained using *HullRad* (Fleming and Fleming 2018) on the initial and optimized ensembles (see **Section S3**).

**Table S4.** Number of residue contacts per conformer between 4E-BP2 and eIF4E for different intermolecular contact regions[a]

| | Overall | RN1 | RN2 | R1 | R2 | R3 | R4 | R5 |
|---|---|---|---|---|---|---|---|---|
| **Initial Pool** | 21.7 | 0.096 | 0.078 | 0.447 | 4.92 | 1.69 | 2.36 | 2.84 |
| **Optimal Ensemble** | 29.6 | 1.69 | 0.35 | 0.85 | 7.49 | 1.49 | 2.70 | 3.74 |
| **Cluster 1** | 21 | 1.5 | 0 | 0.833 | 5.67 | 0.167 | 0.333 | 4.50 |
| **Cluster 2** | 26.1 | 2.37 | 0.184 | 0.947 | 6.71 | 1.13 | 1.76 | 3.24 |
| **Cluster 3** | 35.3 | 1.17 | 0.500 | 0.733 | 8.87 | 2.07 | 3.70 | 4.23 |
| **Cluster 4** | 30.1 | 1.35 | 0.500 | 0.846 | 7.46 | 1.65 | 3.46 | 3.73 |

[a] Regions *R1-R5, RN1 and RN2* as shown in **Fig. 6B** in the manuscript. Contacts were computed for the initial and the optimized 4E-BP2: eIF4E ensemble, in the latter case both as whole and for *Clusters 1-4* resulted from hierarchical clustering.

### REFERENCES


Dawson, J. E., A. Bah, Z. Zhang, R. M. Vernon, H. Lin, P. A. Chong, M. Vanama, N. Sonenberg, C. C. Gradinaru and J. D. Forman-Kay (2020). "Non-cooperative 4E-BP2 folding with exchange between eIF4E-binding and binding-incompatible states tunes cap-dependent translation inhibition." Nat Commun **11**(1): 3146.

Fleming, P. J. and K. G. Fleming (2018). "HullRad: Fast calculations of folded and disordered protein and nucleic acid hydrodynamic properties." Biophysical journal **114**(4): 856-869.

Holehouse, A. S., R. K. Das, J. N. Ahad, M. O. Richardson and R. V. Pappu (2017). "CIDER: resources to analyze sequence-ensemble relationships of intrinsically disordered proteins." Biophysical journal **112**(1): 16-21.

Jain, A., R. Liu, Y. K. Xiang and T. Ha (2012). "Single-molecule pull-down for studying protein interactions." Nature protocols **7**(3): 445-452.

Lee, N. K., A. N. Kapanidis, Y. Wang, X. Michalet, J. Mukhopadhyay, R. H. Ebright and S. Weiss (2005). "Accurate FRET measurements within single diffusing biomolecules using alternating-laser excitation." Biophysical journal **88**(4): 2939-2953.

Ortega, A., D. Amorós and J. G. De La Torre (2011). "Prediction of hydrodynamic and other solution properties of rigid proteins from atomic-and residue-level models." Biophysical journal **101**(4): 892-898.

Shivnaraine, R. V., D. D. Fernandes, H. Ji, Y. Li, B. Kelly, Z. Zhang, Y. R. Han, F. Huang, K. S. Sankar and D. N. Dubins (2016). "Single-molecule analysis of the supramolecular organization of the M2 muscarinic receptor and the Gαi1 protein." Journal of the American Chemical Society **138**(36): 11583-11598.

Smyth, S., Z. Zhang, A. Bah, T. E. Tsangaris, J. Dawson, J. D. Forman-Kay and C. C. Gradinaru (2022). "Multisite phosphorylation and binding alter conformational dynamics of the 4E-BP2 protein." Biophys J **121**(16): 3049-3060.

Tsangaris, T. E., S. Smyth, G.-N. W. Gomes, Z. H. Liu, M. Milchberg, A. Bah, G. A. Wasney, J. D. Forman-Kay and C. C. Gradinaru (2023). "Delineating Structural Propensities of the 4E-BP2 Protein via Integrative Modeling and Clustering." The Journal of Physical Chemistry B **127**(34): 7472-7486.
